# A narrow thermodynamic design window governs selective membrane permeabilization and antiviral activity of amphipathic peptides

**DOI:** 10.64898/2026.04.20.719566

**Authors:** Niek van Hilten, Pascal von Maltitz, Dennis Aschmann, Maria Hoernke, Maximilian Krebs, Jeroen Methorst, Alexander Kros, Jan Münch, Herre Jelger Risselada

**Affiliations:** Leiden Institute of Chemistry, Leiden University, Leiden, The Netherlands; Department of Pharmaceutical Chemistry, Cardiovascular Research Institute, University of California, San Francisco, San Francisco, United States of America; Institute of Molecular Virology, Ulm University Medical Center, Ulm, Germany; Van ‘t Hoff Institute for Molecular Sciences, University of Amsterdam, Amsterdam, The Netherlands; Pharmaceutical Technology and Biopharmacy, Institute of Pharmaceutical Sciences, University of Freiburg, Freiburg, Germany; Physical Chemistry, Martin-Luther-University, Halle (S.), Germany; Department of Physics, Technical University Dortmund, Dortmund, Germany

**Author notes:** These authors contributed equally to this work (Niek van Hilten, Pascal von Maltitz).

## Abstract

Designing molecules that selectively target therapeutically relevant membranes, such as viral envelopes, while sparing host cells is challenging: these membranes closely resemble host bilayers, so selectivity must exploit subtle lipid composition and curvature differences and demands precise tuning of affinity and hydrophobicity, yet curated sequence–specificity data are scarce. Here we show that selective membrane permeabilization and membrane-selective activity of amphipathic peptides are governed by a narrow thermodynamic design window defined by membrane curvature affinity and molecular hydrophobicity. Using a physics-driven generative workflow combining evolutionary molecular dynamics and a transformer predictor (PMIpred), we systematically explored and thermodynamically mapped peptide sequence space *de novo* without reliance on natural templates or experimental training data. Across four design generations we synthesized and experimentally characterized 43 peptides. Mapping functional activity onto a low-dimensional free-energy landscape reveals a confined thermodynamic “sweet spot” separating weak membrane binding from excessive hydrophobic association and cytotoxicity. Peptides operating within this regime efficiently permeabilize model membranes while maintaining low cellular toxicity. Antiviral activity against Zika virus and HIV-1 emerges in the same region but depends sensitively on membrane lipid composition. Quantitative thermodynamic design rules emerge for membrane-active peptides, illustrating how low-dimensional free-energy landscapes can guide the engineering of selective interactions at soft-matter interfaces.

## Introduction

Viruses can be broadly classified into non-enveloped (naked) viruses, which consist solely of a protein capsid enclosing the viral genome, and enveloped viruses, which are surrounded by a lipid bilayer derived from the host-cell membrane during viral budding. Whereas naked viruses rely primarily on capsid-mediated entry mechanisms, enveloped viruses depend critically on the integrity of their lipid envelope for host-cell attachment, membrane fusion, and genome delivery. Disruption of the viral membrane therefore renders enveloped viruses non-infectious. Importantly, many of the most clinically significant human pathogens are enveloped viruses, including Human Immunodeficiency Virus (HIV-1), Influenza virus, and Hepatitis C virus (HCV), as well as emerging viruses from distinct families, such as flaviviruses (e.g., Dengue virus and Zika virus (ZIKV)) and coronaviruses.

To combat viral infection, many therapeutic strategies have been developed to disrupt various stages of the viral reproduction cycle [1], most of which target viral envelope glycoproteins that mediate host-cell recognition and membrane fusion [2–6]. Although mechanistically effective, these proteins exhibit high sequence variability across strains and species [7], facilitating immune escape and limiting cross-protective efficacy. In contrast, the physical properties of viral lipid envelopes—such as curvature, packing defects, and lipid organization—are constrained by their host-derived membrane origin, geometry, and membrane biophysics rather than by genetic sequence. These collective membrane properties therefore provide a conserved physical target space that can be exploited by membrane-active molecules, suggesting that antiviral activity may be governed by thermodynamic interactions with curved lipid membranes rather than by molecular recognition of viral proteins.

Enveloped viruses are nanoscale vesicles typically ranging from 40 to 150 nm in diameter. Clinically relevant flaviviruses such as Dengue virus, Yellow Fever virus, West Nile virus, and Zika virus possess diameters of approximately 40–60 nm (Fig. 1A), i.e. 20–30 nm radii. HIV-1 virions are larger, with diameters near 100 nm. At these dimensions, the positive curvature of the outer membrane leaflet induces lateral strain and increases the density of transient hydrophobic lipid packing defects exposed to the aqueous phase (Fig. 1B). These defects represent thermodynamically unfavorable regions where lipid tails are insufficiently shielded from solvent.

**Fig. 1.**
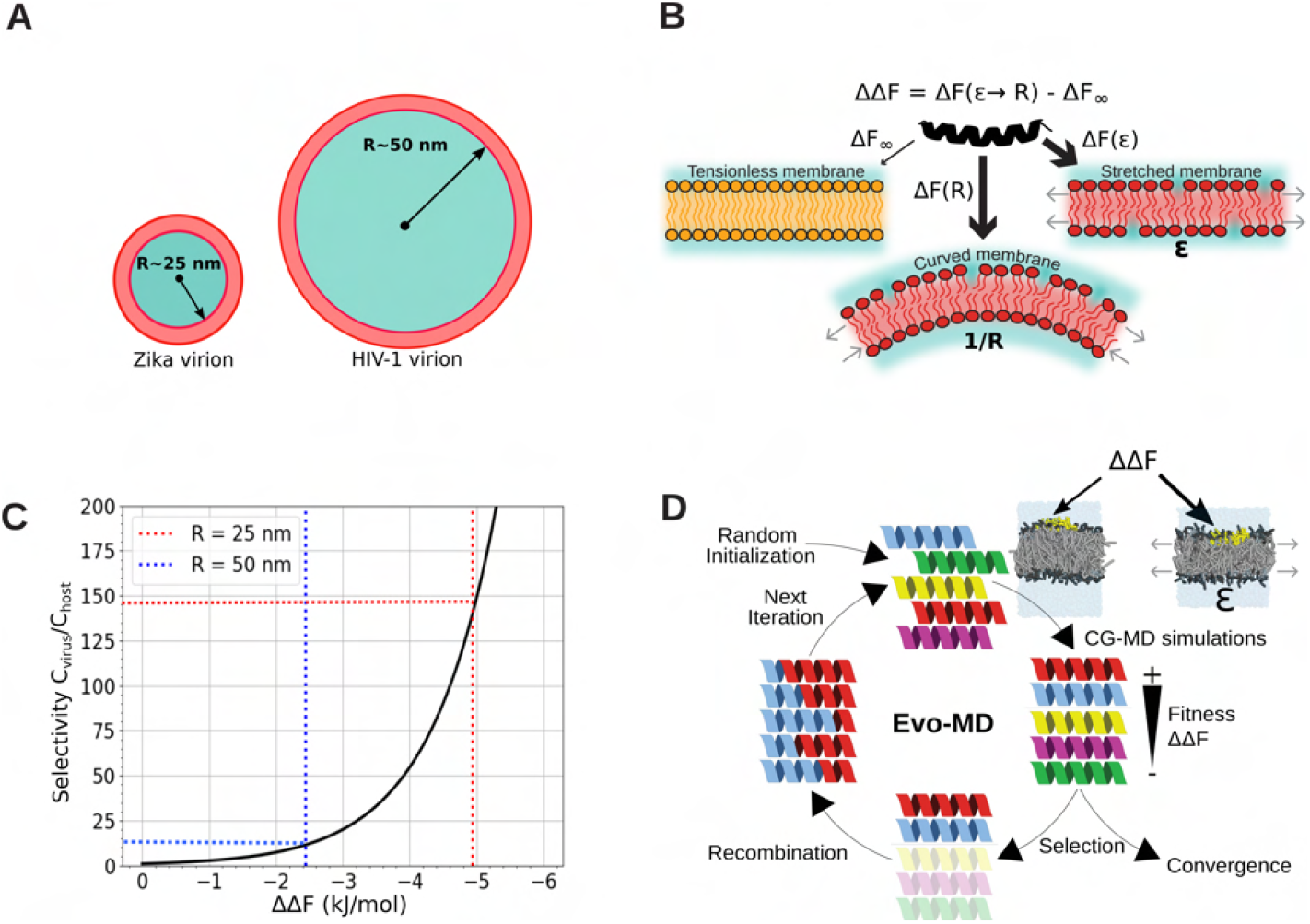
A physics-based generative framework for thermodynamic design of membrane-active peptides. (A) Cartoon representation of the two main viruses targeted by designed peptide drugs and their respective size differences (Figure adapted from Ref. [27]). (B) Schematic representation of curvature-dependent membrane binding. The increased density in lipid packing defects characteristic for positive leaflet curvature (1*/R*) is modeled by a membrane under the relative strain, *ϵ* > 0. The reference binding free energy for a tension-less, planar leaflet (host membrane) is indicated as Δ*F*_∞_. Figure adapted from Ref. [9]. (C) The relative binding free energy ΔΔ*F* determines the ratio between peptide concentration between a vesicle and flat host membrane (coined ‘selectivity’) for the two different virions at thermodynamic equilibrium. The dotted lines indicate the estimated upper binding limit at which peptides demonstrate curvature-sensitive binding before entering a strong-binding regime, for a 25-nm-radius (blue line) and a 50-nm-radius (red line) vesicle, respectively [9, 25]. (D) Exploration of curvature-dependent binding free energies using evolutionary molecular dynamics (Evo-MD) simulations. The peptide within the coarse-grained molecular dynamics simulation system is shown in yellow. The evolution commences with a large pool of randomly generated peptide sequences, simulated in parallel, and progresses based on the peptide’s capacity to preferentially bind a membrane enhanced in hydrophobic packing defects as a result of an externally applied membrane tension, mimicking the effect of positive membrane curvature [24]. The affinity for positive leaflet curvature, ΔΔ*F* is measured from the differential affinity between a small tensionless POPC membrane and a membrane at a relative strain of *ϵ* = 0.165, modeling a vesicle with a radius of *R* ≃ 12.5 nm. Figure adapted from Ref. [9].

Amphipathic peptides can exploit such packing defects by inserting their hydrophobic face into exposed regions while maintaining polar contacts with water, a phenomenon known as lipid packing defect sensing [8–11]. Natural curvature-sensitive motifs, including Amphipathic Lipid Packing Sensor (ALPS) domains, have been shown to preferentially associate with positively curved membranes [12–15]. These findings suggest that curvature-dependent membrane recognition can be described within a thermodynamic framework and may be harnessed for molecular design.

A compelling biological precedent for membrane-targeting antiviral activity is provided by amphipathic helices derived from the hepatitis C virus NS5A protein. The NS5A amphipathic helix (HCV AH) and its overlapping derivative C5A combine low toxicity with broad-spectrum antiviral activity against enveloped viruses, including ZIKV and HIV-1 [16–23]. These peptides disrupt viral envelopes through membrane-permeabilizing mechanisms. In liposomal assays, HCV AH displays vesicle-size–dependent permeabilization, consistent with curvature-sensitive binding [17, 18, 22], whereas C5A exhibits stronger overall membrane binding and reduced curvature discrimination [19, 22]. These peptides demonstrate that therapeutic potential – antiviral activity coupled with low toxicity – depends on a balance of curvature preference, hydrophobicity, and membrane affinity rather than on curvature selectivity alone.

The affinity of a peptide for lipid packing defects (ΔΔ*F*) is thermodynamically characterized by the difference in binding free energy between a flat and tension-free membrane (Δ*F*_∞_) and a membrane with positive curvature (Δ*F*(*R*)), which can alternatively be described as the affinity for a stretched membrane leaflet (Δ*F*(*ϵ*)) [24]. This thermodynamic relationship is sketched in Fig. 1B.

In thermodynamic equilibrium, the selectivity for a membrane is characterized by the ratio of peptide concentrations between the viral membrane (*C*_*virus*_) and the host membrane (*C*_*host*_), as well as the damage inflicted. This binding selectivity is quantified by the expression 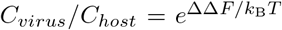, where *k*_B_ represents Boltzmann’s constant and *T* denotes the absolute temperature in Kelvin. This equation indicates that the relative concentration of peptides on the viral membrane, compared to the host membrane, varies exponentially with changes in the relative membrane affinity (ΔΔ*F*, see Fig. 1C).

Consequently, a peptide with an affinity for positive leaflet curvature (ΔΔ*F* < 0) is expected to have a significantly higher concentration on the membrane surface of small virions than on the host membrane. The more negative the value of ΔΔ*F*, the larger the peptide’s affinity for positive curvature. For 25-nm-sized vesicles (*R* = 12.5 nm), the standard reference size used in our Protein Membrane Interaction prediction (PMIpred) server [25], a curvature-sensing sequence has a relative binding free energy (ΔΔ*F*) of approximately 10 kJ/mol compared to a flat, tensionless membrane [9]. This value decreases with increasing vesicle size, and its scaling with liposome radius can be estimated using a previously published relationship (see Ref. [10]) [9, 25]. Consequently, targeting a Zika virus particle (*R* = 25 nm) by curvature-sensing sequences is estimated to lead to a 150-fold increase in peptide concentration on the viral membrane compared to the host membrane (Fig. 1C). Similarly, targeting a larger HIV-1 virion (*R* = 50 nm) results in a roughly 12-fold increase in peptide concentration on the viral membrane (Fig. 1C).

Our previous work demonstrated that an amphipathic peptide’s preference for hydrophobic lipid packing defects correlates directly with its capacity to generate leaflet stress [24], establishing a thermodynamic connection between curvature affinity and membrane permeabilization. Yet strengthening curvature affinity through increased hydrophobicity also enhances nonspecific membrane association, self-aggregation, and cytotoxicity. Functional performance is therefore confined to a bounded region of thermodynamic space rather than scaling monotonically with ΔΔ*F*.

Despite the therapeutic potential of selective membrane-targeting antiviral peptides (AVPs), experimentally validated sequence data remain limited, restricting purely data-driven generative approaches [26]. To address this gap, we developed a physics-driven framework combining evolutionary algorithms with coarse-grained molecular dynamics (Evo-MD) to generate peptides directly from curvature-dependent binding free energies [9]. As demonstrated in Ref. [9], integrating Evo-MD with a transformer-based membrane interaction predictor and statistical mechanical modeling successfully recovered known ALPS motifs and their mutants without relying on training data.

We extended this Evo-MD framework to enable bottom-up, physics-driven design of membrane-targeting antiviral peptides. A detailed characterization of the lead candidate from this effort, P1.6, was recently reported [23]. P1.6 inhibits ZIKV and HIV-1 infection at sub-micromolar concentrations, matching or surpassing established amphipathic antiviral peptides, while exhibiting minimal cytotoxicity and no detectable hemolytic activity [23]. In the current work, we systematically refine the generative framework across four iterative design generations. By tuning algorithmic constraints and exploring defined regions of curvature affinity and hydrophobicity space, we designed and experimentally evaluated 43 *de novo* peptide sequences (Table SI1). Five candidates demonstrated antiviral activity against ZIKV and/or HIV-1 (Table 1). Integration of our dataset with known membrane-active sequences from the literature reveals that antiviral efficacy emerges from a balance between curvature-dependent membrane affinity, sufficient hydrophobicity to induce membrane stress, and avoidance of peptide self-aggregation. Notably, P1.6 exhibits potent viral membrane disruption in transmission electron microscopy (TEM), yet does not display curvature-sensitive permeabilization in standard liposome leakage assays. This observation indicates that antiviral selectivity does not necessarily require strict curvature sensing in simplified model systems [22], but instead arises from operating within an optimal thermodynamic regime that balances binding strength and leaflet stress generation.

**Table 1:** Overview of properties for generated sequences that showed antiviral activity. Summary of viral inhibition, cell viability, and hemolysis results for active peptides.

| | $\Delta\Delta F_{adj}$ (kJ mol<sup>-1</sup>)<br>PMIPred | $\Delta F_{\infty}$ (kJ mol<sup>-1</sup>)<br>interpolation/simulation | $\langle H \rangle$ | ZIKV | | | HIV-1 | | | Hemolytic |
| --- | --- | --- | --- | --- | --- | --- | --- | --- | --- | --- |
|  |  |  |  | IC<sub>50</sub> | CC<sub>50</sub> | SI | IC<sub>50</sub> | CC<sub>50</sub> | SI |  |
| DMF |  |  |  | X | >20 | X | X | >20 | X | No |
| HCV AH | -11.5 | -31.8 / -16 ± 5 | 0.78 | 1.2<sup>†</sup> | >20 | >16.7 | 0.1<sup>†</sup> | >20 | >200 | No |
| C5A | -15.0 | -45.0 / -18 ± 9 | 0.74 | 0.8<sup>†</sup> | >20 | >20 | 0.1<sup>†</sup> | >20 | >200 | Yes |
| P1.6 | -10.5 | -28.0 / -18 ± 3 | 0.77 | 0.3<sup>†</sup> | >20 | >71.4 | 2.5<sup>†</sup> | >20 | >8.0 | No |
| P2.2* | -30.4 | -104.1 / -53 ± 3 | 1.0 | X | >20 | X | X | 6.8 | X | No |
| P2.2.3 | -22.5 | -73.9 / -22 ± 2 | 0.6 | 1.1 | 8.6 | 8.0 | 1.2 | 6.3 | 5.2 | Yes |
| P2.2.4 | -25.3 | -84.6 / -23 ± 5 | 0.6 | 1.3 | 2.9 | 2.3 | 0.4 | 5.1 | 14.2 | Yes |
| P2.2.4.d | -23.0 | -75.8 / -33 ± 8 | 0.6 | 3.6 | 10.0 | 2.8 | X | 10.5 | X | No |
| P2.4.d | -21.3 | -65.8 / -19 ± 7 | 0.57 | 1.4 | >20 | >14.2 | X | 17.9 | X | No |

Together, our results reveal a narrow thermodynamic design window governing selective membrane permeabilization by amphipathic peptides, establishing quantitative design rules for membrane-targeting antiviral molecules.

## 1 Results

### Physics-driven exploration of the thermodynamic design landscape

Our central hypothesis is that selective membrane permeabilization by amphipathic peptides emerges only within a confined thermodynamic regime defined by curvature affinity and hydrophobicity. To identify this regime, we used a physics-driven generative design strategy that systematically explores peptide sequence space based on curvature-dependent membrane binding free energies. This approach integrates evolutionary molecular dynamics simulations (Evo-MD) with our transformer-based Protein–Membrane Interaction predictor (PMIpred), enabling efficient navigation of the thermodynamic landscape governing peptide–membrane interactions (Fig. 1D). By iteratively generating and experimentally testing candidate sequences across four design generations, we mapped how curvature affinity and hydrophobicity jointly determine membrane permeabilization, antiviral activity and cytocompatibility.

Evo-MD combines a Darwinian evolutionary algorithm with coarse-grained molecular dynamics simulations to generate peptide sequences optimized for curvature-dependent membrane binding. Candidate sequences are evaluated using a fitness function derived from ensemble-averaged simulation trajectories that quantify the relative binding free energy ΔΔ*F*, representing the affinity of a peptide for lipid packing defects associated with positive membrane curvature (Fig. 1A,B). Because sequence generation is guided entirely by physical interaction energies, the approach requires no experimental training data and can explore peptide sequence space in a fully physics-based manner [9]. Across four design generations we systematically sampled distinct regions of this thermodynamic landscape to determine where selective membrane permeabilization and antiviral activity co-localize.

The vast size of the search space (20^*N*^ possible peptide sequences for 20 natural amino acids and sequence length *N*) necessitates efficient sampling strategies. We therefore implemented a Darwinian evolutionary algorithm that iteratively evaluates candidate fitness and generates new variants via mutation and recombination (Fig. 1D). A sequence length of *N* = 24 was chosen to approximate bilayer thickness (approximately 4 nm), consistent with toroidal pore formation as a membrane-permeabilizing mechanism [16]. Simulations were performed using the Martini 3 force field [28–30] to enable high-throughput exploration.

Within this framework, ΔΔ*F* serves as the central thermodynamic descriptor. It approximates the difference in binding affinity between a stretched membrane (Δ*F*(*ϵ*)) and a flat membrane (Δ*F*_∞_) [24]. Increasingly negative ΔΔ*F* values correspond to stronger membrane association and enhanced leaflet stress generation. However, strong membrane binding also promotes reduced reversibility and increased nonspecific interactions, indicating that functional activity is confined to a bounded thermodynamic region rather than increasing monotonically with ΔΔ*F*.

Previously, we demonstrated that sequences driven toward extreme curvature affinity converge to highly hydrophobic peptides with limited aqueous solubility [9]. To preserve biological relevance, we constrained subsequent exploration to sequences predicted to remain soluble and helical (Table SI1).

In total, 43 *de novo* peptides were generated and experimentally characterized across four generations (P1–P4), each validated in antiviral and liposome leakage assays to map functional outcomes onto the thermodynamic landscape.

All four generations sampled complementary regions of this design space:

- P1.x-force (9 sequences): Curvature-dependent binding was initially explored using a sorting-force–based metric [10]. This generation yielded the lead sequence P1.6, which potently inhibits HIV-1 and ZIKV while remaining non-hemolytic (Table 1, Fig. 2A) [23].
- P2.x-constraint (14 sequences): Sequences with progressively stronger curvature affinity were examined under solubility constraints. Several displayed potent membrane permeabilization and significant antiviral activity but increased cytotoxicity, consistent with occupation of the strong-binding region of the landscape.
- P3.x-maxsol (10 sequences): The low-hydrophobicity region was probed via multi-objective exploration of curvature affinity and mean hydrophobicity [31]. These peptides showed excellent solubility but limited membrane permeabilization and lacked antiviral activity, indicating insufficient membrane association.
- P4.x-sweetspot (10 sequences): Sequences positioned within the intermediate curvature–hydrophobicity regime exhibited strong membrane permeabilization with low cytotoxicity. However, the absence of concurrent antiviral activity exposed the limits of the thermodynamic design window, revealing dependencies on additional features of the viral membrane composition.

**Fig. 2.**
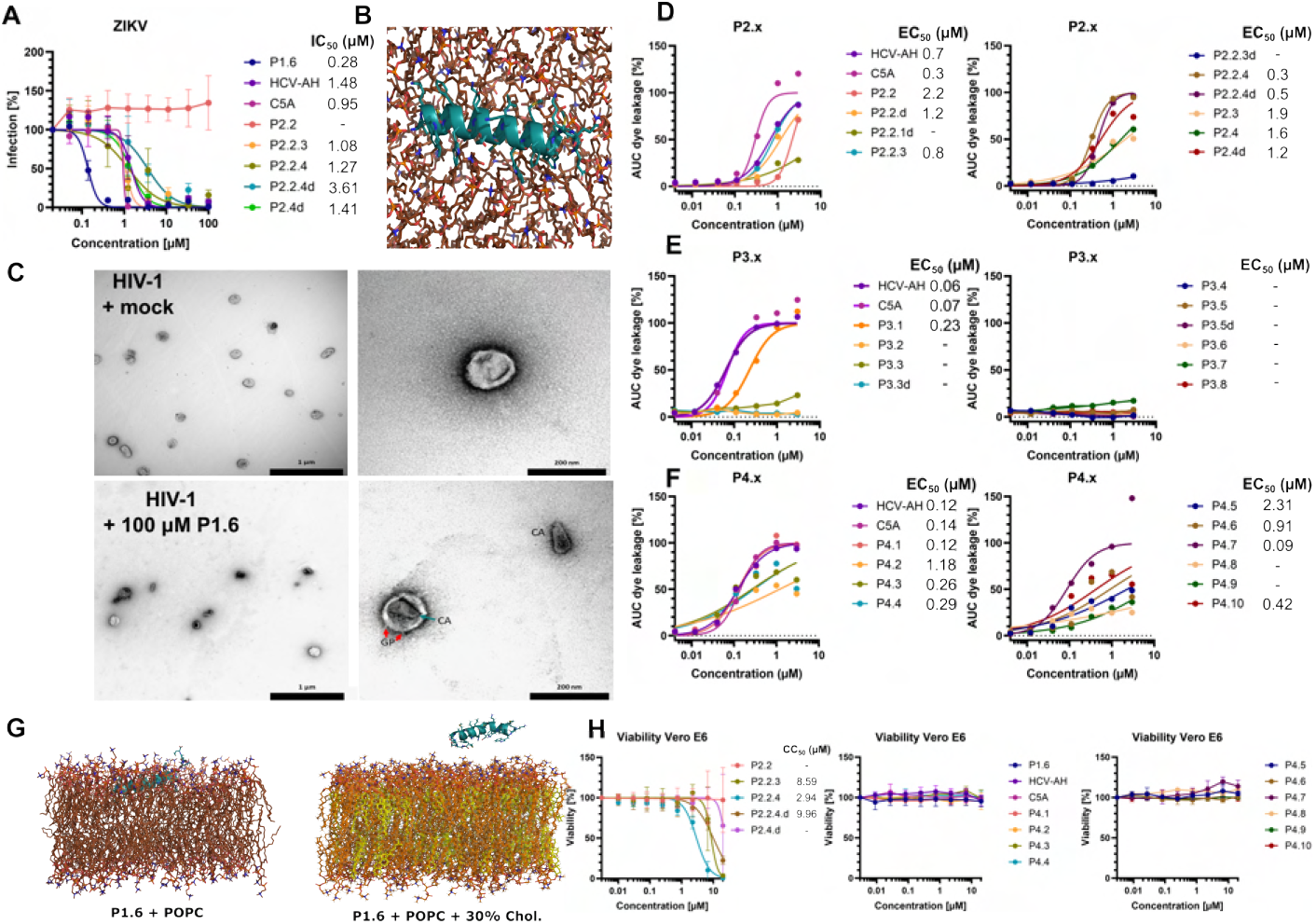
Experimental validation of selective membrane permeabilization and antiviral activity across the thermodynamic design landscape. (A) ZIKV infection assays on Vero E6 cells following incubation with different peptide concentrations. Shown are all peptide candidates developed in this study that exhibited antiviral activity, including known active controls (HCV AH and C5A). (B) Atomistic simulation of the lead sequence P1.6 (top view) bound to a POPC membrane (Adapted from Ref. [23]). (C) TEM images of HIV-1 virions incubated with either peptide P1.6 or with vehicle only (mock-treated control). Peptide P1.6 removes the viral lipid envelope, leaving a bare capsid (CA).(D–E) Liposome dye leakage assays using 100 nm liposomes (The lipid composition of 45/25/30 mol% DOPC/SM/cholesterol conforms to Ref. [32].) for generations P2.x, P3.x, and P4.x. All sequences from generation P4.x were inactive in viral assays despite featuring potent membrane-permeabilizing activity against liposomes. (G) Atomistic molecular simulation illustrating how the addition of 30% cholesterol to a POPC membrane impairs membrane binding of the lead sequence P1.6 (Adapted from Ref. [23]). (H) Vero E6 cell viability upon exposure to varying peptide concentrations, comparing the generations P2.x and P4.x plus controls. Virus assays (N=3), viability assays (N=2), and liposome leakage assays (N=1) were each performed in triplicate.

Extensive details on the four design strategies and remaining data are supplied in the Supplemental Methods and Supplemental Information. In the main manuscript, we will predominantly focus on the key insights that followed from integrating all the gathered data.

To provide an insightful overview of the total dataset, we visualized the combined data by mapping the peptides’ effective affinity for positive leaflet curvature against the peptides’ mean hydrophobicity (Fig. 4A). Active AVPs are indicated with a star symbol (⋆) and inactive peptides with a circle symbol (⋆). Overall, our analysis encompasses more than 50 sequences, including our own experimentally validated designs (Table SI1), along with known curvature-sensing sequences [9] and known antiviral sequences from the literature [16, 19, 33, 34].

Two descriptors characterize peptide activity:

- *Adjusted Free Energy (*ΔΔ*F*_*adj*_). The here-reported ΔΔ*F*_*adj*_ values are predicted by our PMIpred transformer model that was trained on Evo-MD data for > 50, 000 peptide sequences [25]. The adjusted free energy enables a fair comparison between different peptides by applying a fixed energy scale corresponding to a vesicle radius of 12.5 nm. This scale is conform to the original stressed membrane setup at a relative strain of *ϵ*_*ref*_ = 0.165 at which the Evo-MD simulations were performed [9, 24]. ΔΔ*F*_*adj*_ also adjusts for variations in peptide length and net charge [9]. All reported values align with predictions made within our publicly accessible PMIpred server [25].
- *Mean Hydrophobicity (*⟨ *H* ⟩) The mean hydrophobicity of a peptide sequence ⟨ *H* ⟩ is determined by summing the hydrophobicity values of each amino acid in the sequence and then dividing by the total number of amino acids:

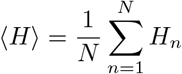

This calculation aligns with the procedure outlined on the HeliQuest web-server [35], which uses hydrophobicity values derived from the octanol/water partition coefficient [36]. Higher values indicate greater hydrophobicity.

Projecting peptide activity onto this two-dimensional descriptor space reveals a confined thermodynamic “sweet spot” where peptides are prone to bind to highly curved membranes (ΔΔ*F*_*adj*_ < −10 kJ mol^−1^), while having a balanced hydrophobicity (0.5 ≤ ⟨ *H* ⟩ ≤ 1.0). The four design generations progressively delineate a narrow thermodynamic design window in which selective membrane permeabilization and antiviral activity co-localize.

### Generation 1. Potent membrane-permeabilizing antivirals emerge near the weak-binding boundary

Previously, we illustrated that a peptide’s affinity for positive membrane curvature is equivalent to the peptide’s propensity for exerting leaflet stress [24]. This highlights how curvature-dependent interactions shape the thermodynamic landscape governing membrane-permeabilizing activity. Additionally, we demonstrated that the peptide’s preference for positive leaflet curvature, ΔΔ*F* correlates with its affinity for non-curved membranes, Δ*F*_∞_ (Fig. 4B and Fig. SM5). This ‘sensing paradox’ arises because both phenomena are driven by hydrophobicity. That is, an increased curvature preference paradoxically enhances binding across all degrees of membrane curvature — including flat membranes — ultimately leading to toxic effects on the host cell.

Only within the narrow curvature-sensing regime (Fig. 4C) do peptides discriminate between curved and non-curved membranes, as membrane association remains reversible on biologically relevant timescales – an essential prerequisite for curvature sampling. Outside this regime, the binding probability approaches unity irrespective of vesicle radius (Fig. 4D), and the slower unbinding kinetics lead to effectively irreversible attachment, eliminating curvature selectivity.

As previously noted, peptides beyond the curvature sensing range generate excessive stress in the outer leaflet, leading to enhanced membrane-permeabilizing activity [9, 24]. Antiviral performance emerges from a balance between membrane association strength and cytocompatibility within this thermodynamic landscape. Small enveloped viruses, particularly flaviviruses (*R* ≈ 25 nm) like Dengue, Yellow Fever, and Zika viruses, are expected to be most vulnerable to curvature-biased thermodynamic enrichment (Fig. 1C).

This raises the question of where functional membrane-permeabilizing activity resides within the thermodynamic landscape. Curvature-sensing peptides, exemplified by ALPS motifs [13], typically exhibit adjusted binding free energies (ΔΔ*F*_*adj*_) in the range of 6 to 10 kJ mol^−1^, as estimated using the PMIpred metric [9, 25]. Thus, this regime corresponds to the weak yet selective membrane binding observed in standard liposomal assays, which typically employ 100 nm-sized liposomes and peptide concentrations around 1 mM. It serves as a criterion for classifying curvature sensing within our PMIpred server [25] (Fig. 4C). Beyond this range, peptides are expected to bind strongly to liposomes regardless of their size (binding probability approaching unity), whereas below this range they remain unbound in solution (binding probability approaching zero). Two of the most effective AVPs against Zika virus, HCV AH and P1.6, exhibited a relative binding free energy slightly above the estimated upper threshold for curvature sensing, at about ΔΔ*F*_*adj*_ = 10 kJ mol^−1^(Table 1 and Fig. 4A).

In parallel to our theoretical analysis and antiviral assays, we conducted liposome leakage experiments. These measurements revealed no size-dependent variation in the membrane-permeabilizing activity of P1.6 or the other active peptides from generation P2.x across vesicles of 50, 100, and 200 nm diameter (see Ref. [23], Table SI3, Fig. SI12). This behavior mirrors the well-characterized mechanism of the antiviral peptide C5A [22], yet contrasts sharply with HCV AH, whose permeabilizing potency increases as liposome size decreases [17, 18, 23, 38]. Strikingly, the ΔΔ*F*_*adj*_ values for P1.6 and HCV AH are nearly identical and very close to the boundary between weak binding (sensing) and strong membrane association (‘weak binding regime’, Fig. 4A,C). Both P1.6 and HCV AH exhibit substantially higher hydrophobicity than most native curvature-sensing motifs (dark blue closed circles in Fig. 4A), which we propose may increase their permeabilizing potency. Indeed, our dataset indicates that effective membrane-permeabilizing peptide sequences require an average hydrophobicity of at least 0.5. Logically, such strong membrane-permeabilizing properties are undesirable [39] for native curvature sensors because their interactions with lipid membranes must remain inert and reversible, explaining their generally more hydrophilic character.

Summarizing the findings for P1.6 and state-of-the-art AVPs HCV AH and C5A, we propose that membrane-permeabilizing antiviral peptides achieve optimal performance within the weak binding regime - near or slightly above the curvature-sensing threshold. In this regime, membrane association is sufficiently strong to promote antiviral efficacy without causing extensive cellular membrane damage. Although HCV AH exhibits pronounced vesicle-size-dependent permeabilizing activity, its curvature-dependent binding appears secondary to the thermodynamic balance underlying viral selectivity. From a thermodynamic perspective, increasing binding probability enhances leaflet tension and thereby promotes permeabilizing activity. When membrane binding becomes effectively irreversible on biologically relevant timescales, true curvature sensing is lost; however, selectivity may then depend on cellular membrane repair mechanisms. Such processes are absent in enveloped viruses. Consequently, optimal antiviral binding likely occurs within a regime where biophysical assays no longer detect curvature-sensing behavior.

### Generation 2. More is less: Increasing curvature affinity beyond the optimal window reduces antiviral selectivity

Generation P2.x systematically probed the high-curvature-affinity region of the thermodynamic landscape while constraining peptide solubility by fixing the number of hydrophilic residues to a predefined fixed number. In total, 10 sequences were selected with ΔΔ*F*_*adj*_ values ranging from about -15 to -34 kJ mol^−1^, considerably higher curvature affinities than those of HCV AH and P1.6. Four sequences, P2.2.3, P2.2.4, P2.2.4 d, and P2.4 d showed moderate (≤ 4 *µ*M) to good (≤ 1 *µ*M) IC_50_ activity in viral infectivity assays against ZIKV and/or HIV-1 (Table 1). All four were highly active in liposome dye leakage assays (EC_50_ < 2 *µ*M), confirming membrane-permeabilizing capability (Table SI3, Fig. 2D). Overall, generation P2.x displayed strong membrane-permeabilizing capabilities, probably explained by the relationship between curvature affinity and leaflet stress generation [9, 24]: high curvature affinity apparently leads to high leaflet stress and thus greater propensity for membrane permeabilization. However, as high curvature affinity also correlates with strong general membrane binding of these peptides, they performed moderately to poorly in hemolysis and cell viability assays compared to HCV AH and P1.6 (Table 1, Fig. 2H, Fig. SI11). These toxic effects may limit the clinical potential to topical prophylaxis against viral infection, similar to C5A [19, 40], used as a positive control here. Notably, C5A and P2.2.3 exhibited similar activity against ZIKV, with IC_50_ values of about 1 *µ*M (Table 1).

In order to examine the membrane binding characteristics of the here-uncovered active peptide sequences, we performed adsorption experiments on DMPC monolayers (Fig. SI13B). Specifically, we monitored changes in surface pressure (Δ*π*) from the initial value (*π*_0_) upon addition of a peptide solution to the subphase of a DMPC monolayer (see section SM3 for methodological details). We observed that P2.2.3, and not P1.6, associated with the air-water interface in absence of DMPC (*π*_0_ = 0), in line with P2.2.3 being more hydrophobic than P1.6. Moreover, the maximal insertion pressures (*π*_mip_, i.e. the intercept with Δ*π* = 0), for both P2.2.3 and P1.6 are above the DMPC monolayer-bilayer correspondence pressure (30-35 mN m^−1^, [41, 42]), indicating that both peptides would readily insert into PC bilayers (*π*_mip_ > 45 mN m^−1^). In our observations, the non-active control peptide sequence P2.2 failed to integrate into the DMPC monolayer, despite its notably larger hydrophobicity (⟨ *H* ⟩ = 0.957 for P2.2, compared to ⟨ *H* ⟩ = 0.557 for P2.2.3). We assume that this may be due to strong self-interactions between the peptide molecules, causing them to aggregate in solution, and thus preventing the peptides to insert into the lipid monolayer.

Intriguingly, the four active peptides in generation P2.x have a mean hydrophobicity value of around 0.6. All peptides with higher ⟨ *H* ⟩ values were inactive. In fact, some of the peptides, especially the more hydrophobic ones, visibly aggregated during synthesis, which in some cases rendered the peptides insoluble (Table SI1). Although peptides in generation P2.x seem to bind stronger to membranes than the lead antiviral sequence P1.6 and, thus, are expected to generate more leaflet strain, both their membrane-permeabilizing and antiviral activity tend to be lower than that of P1.6.

This leaves us with the question of *why* the four active P2.x peptides were less potent than P1.6, despite their larger (*i*.*e*., more negative) ΔΔ*F*_*adj*_ values. Together with their increased toxicity, we attribute these observations to several factors:

1. Larger negative ΔΔ*F*_*adj*_ values directly reflect the peptides’ propensity to induce more leaflet stress in a pure POPC membrane subject to strong positive membrane curvature [24]. However, our recent atomistic simulations of P1.6 revealed that it binds potently to pure POPC membranes, but not to membranes with 30% cholesterol (see Ref. [23] and Fig. 2G). This additional dependence on lipid composition is evident in the differential activity profiles exhibited by the tested sequences across distinct virus types and liposome compositions [23], thus muddying direct translation from prediction to experiment.
2. Peptides with larger ΔΔ*F*_*adj*_ typically feature a wide, unperturbed, hydrophobic interface (Fig. 3, Fig. SI2). We assume that this promotes self-aggregation and thus reduces the concentration of peptides available for membrane binding. An illustrative example of this phenomenon is P2.2, which, despite its estimated intrinsic strength as a membrane binder (with a Δ*F*_∞_ of > 50 kJ mol^−1^), does not insert into membranes (Fig. SI13B).
3. Larger negative ΔΔ*F*_*adj*_ values are associated with increased non-specific membrane binding (Δ*F*_∞_) [9, 25]. Unbinding through thermal fluctuations therefore might pose a significant challenge within the timescale of cellular activities. Consequently, binding becomes apparently irreversible regardless of curvature. In a situation where membranes of different curvatures ‘compete’ for an AVP, this phenomenon reduces the concentration of peptides available for subsequent viral membrane binding, thereby diminishing antiviral efficacy. Specifically, their ability to non-specifically bind to membranes with high potency increases their potential to cause cellular toxicity.

**Fig. 3.**
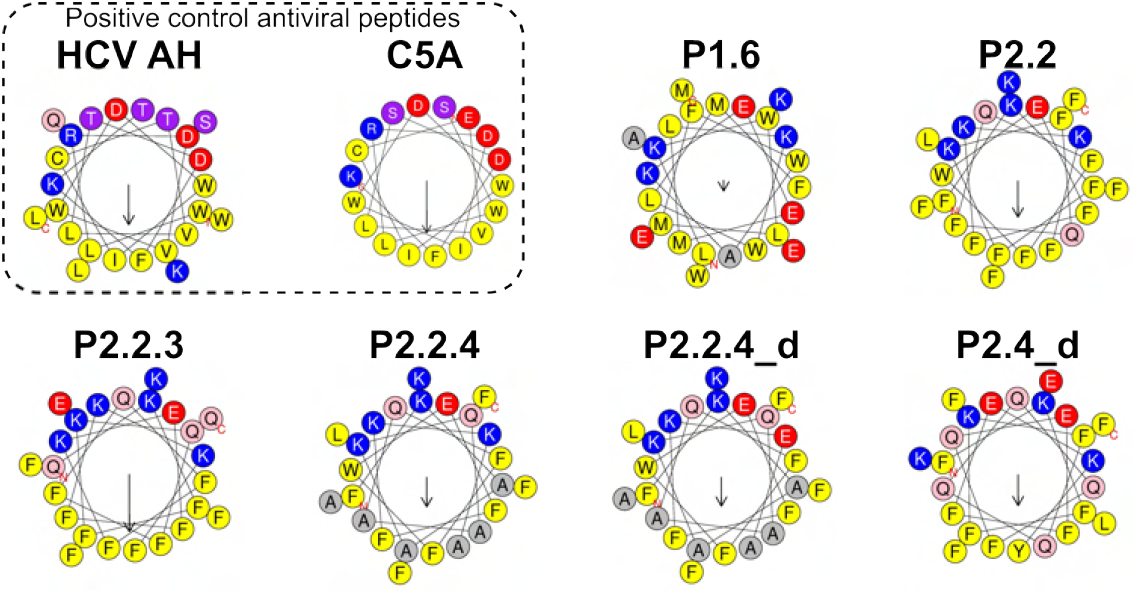
Helical wheel representations of generated sequences showing antiviral activity. Helical wheel projections were generated using HeliQuest [37] for the peptides in Table 1. Helical wheel plots for all other sequences are provided in Fig. SI1, Fig. SI2, Fig. SI3B, and SI5. Arrows indicate both the direction and magnitude of the mean hydrophobic moment.

**Fig. 4.**
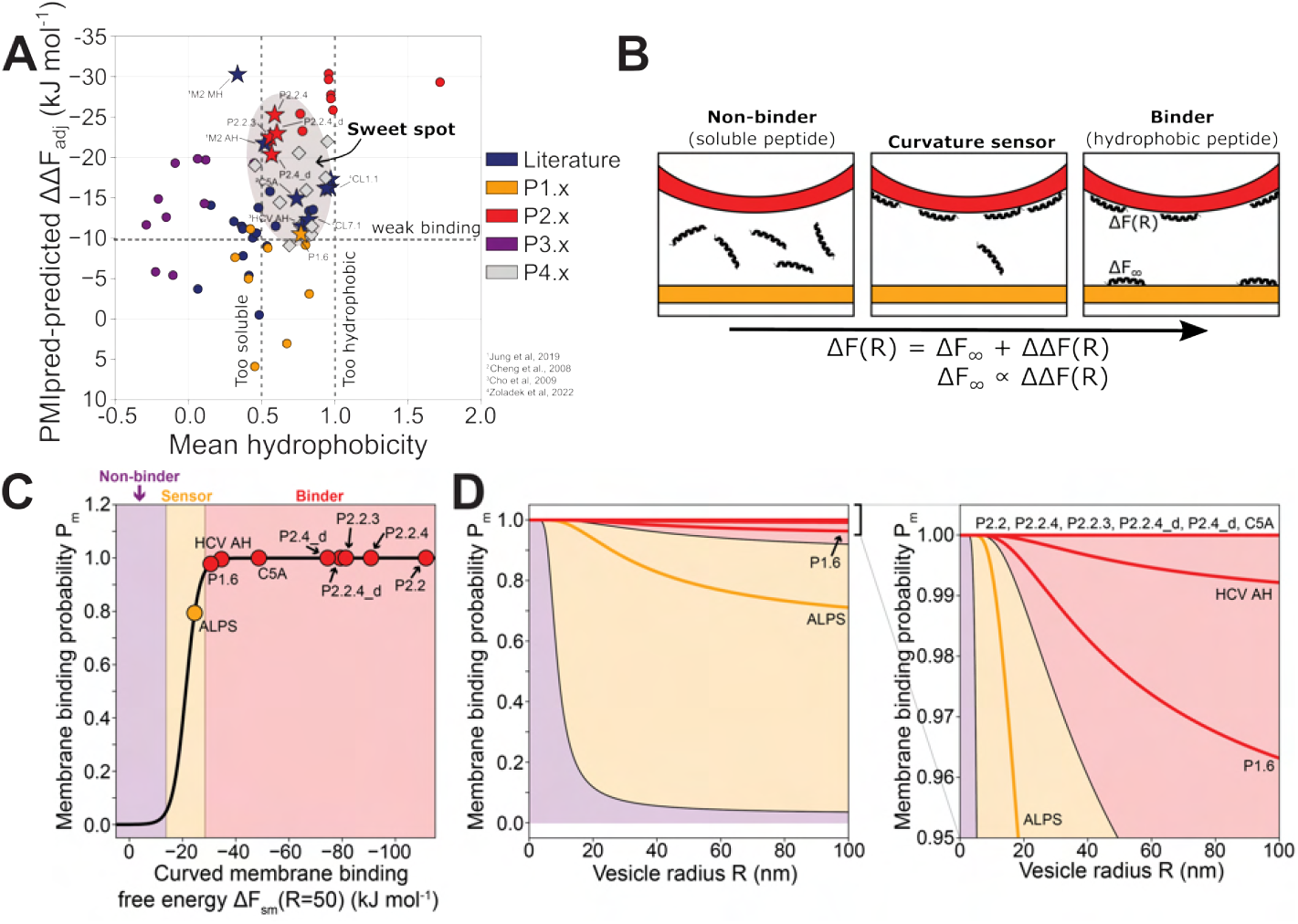
Mapping the thermodynamic design landscape of membrane-active peptides across four generations. (A) We conceptualize antiviral activity through two key peptide descriptors: preference for positive leaflet curvature (ΔΔ*F*_*adj*_), as predicted by our PMIpred server [25], and mean hydrophobicity ⟨ *H* ⟩. Active AVPs are highlighted with a star symbol. (B) Thermodynamics of curvature ‘sensing paradox’: enhancing a peptide’s ability to bind to curved surfaces Δ*F* (*R*) also improves its binding to flat membranes Δ*F*_∞_ because both are driven by hydrophobic interactions [9]. (C) The selective interaction of peptides with small vesicles (curvature sensing) can only occur within a very narrow range of Δ*F* (*R*), or equivalently, a narrow range of ΔΔ*F*_*adj*_ which is approximated to fall within about 6 up to 10 kJ mol^−1^(scaled values) [9, 25]. (D) True curvature sensors like ALPS possess a unique characteristic: their membrane binding probability, specifically the adopted peptide concentration at the membrane, exhibits strong dependence on vesicle size. Peptides in the ‘binder’ regime bind to membranes independent of vesicle size.

In summary, although movement toward more negative ΔΔ*F* values shifts sequences deeper into the strong-binding region of the landscape, their potent membrane binding and concomitant strong membrane-permeabilizing properties actually make them more prone to non-specific binding and hemolysis [39]. Indeed, P2.x peptides typically showed reduced cell viability in our experiments (Fig. 2H).

### Generation 3. A minimal hydrophobicity is required for antiviral activity

One of the main hurdles we faced in generation P1.x and P2.x was peptide solubility. Maximizing ΔΔ*F*_*adj*_ tends to produce highly hydrophobic sequences that are prone to self-aggregate in solution, leading to low antiviral activity.

To overcome these challenges, the third generation (P3.x) was designed with a multi-objective optimization technique. This strategy aimed to *maximize* curvature affinity while simultaneously *minimizing* mean hydrophobicity.

As a result, ten peptide sequences were selected, each with unique balances of ⟨ *H* ⟩ and ΔΔ*F*, identified after evolutionary optimization. As intended, peptides in the P3.x series exhibited superior solubility, with mean hydrophobicity values ranging from -0.3 to 0.4, notably lower than sequences from earlier generations (Table SI1). Additionally, eight of these peptides surpassed the curvature sensing threshold, with adjusted relative binding free energy (ΔΔ*F*_*adj*_) values ranging from -10 to -20 kJ mol^−1^.

Further investigation involved assessing the membrane-permeabilizing activity of these peptides using liposome dye leakage assays [43] (Fig. 2E). Peptide P3.1 - with ⟨ *H* ⟩ = 0.447 the most hydrophobic sequence of its generation (Table SI1) - was the only P3.x peptide that exhibited promising activity in our liposome leakage assays with an IC_50_ of 0.23 *µ*M, approximately four to five fold lower than HVC AH and C5A. Disappointingly, however, P3.1 did not inhibit HIV-1 or ZIKV infections (Fig. SI16 and Fig. SI17), indicating a lack of sufficient potency to disrupt viral membranes. We argue that these peptides were likely *too soluble*. Together, these findings suggest that potent membrane-permeabilizing antiviral peptides typically possess mean hydrophobicity values greater than ≈ 0.5. In fact, while beyond the focus of our study, it is conceivable that some of the P3.x sequences might exhibit behavior akin to native curvature-sensing sequences due to their high solubility and lack of permeabilizing interaction with membranes.

Combining knowledge from the initial three peptide generations indicates that effective potent, membrane-permeabilizing antiviral peptides with low toxicity likely have ΔΔ*F*_*adj*_ values between -10 and -25 kJ mol^−1^and a mean hydrophobicity within a narrow range, slightly above 0.5 but not exceeding 1.0: the AVP sweet spot (Fig. 4A).

### Generation 4. Operating within the thermodynamic sweet spot: selective membrane permeabilization with low cytotoxicity

In the final design round, generation P4.x, we selected 10 peptides positioned within the identified thermodynamic sweet spot from all our previously generated sequences as well as the antiviral sequences reported in the literature [16, 19, 33, 34]. This ‘sweet spot’ in chemical space is highlighted by the gray ellipse in Fig. 4A.

Indeed, the majority of peptides from generation P4.x showed substantial concentration-dependent activity in leakage assays utilizing 100 nm sized model liposomes (Fig. 2F), with low or sub-micromolar *IC*_50_ values for 7 of the 10 peptides. Remarkably, P4.1 and P4.7 exhibited membrane-permeabilizing potencies matching or exceeding those of the control peptides HCV AH (0.12 *µ*M) and C5A (0.14 *µ*M), with IC_50_ values of 0.12 *µ*Mand 0.09 *µ*M, respectively. Furthermore, Vero E6 cell viability assays showed no cytotoxicity for the active P4.x peptides even at the highest tested concentrations of up to 20 *µM* (see Fig. 2H). These results demonstrate that sequences residing within the thermodynamic sweet spot exhibit selective membrane permeabilization with no cytotoxicity.

However, a notable disparity emerged between the peptides’ permeabilizing activity toward virus-like liposomes and their antiviral efficacy: despite exhibiting potent liposome permeabilization, they only showed weak to moderate inhibition in ZIKV infectivity assays (Fig. SI18), with P4.3 having the lowest IC_50_ at 1.8 *µ*M. In a head-to-head comparison, however, it was approximately 5 and 90 times weaker than C5A and HCV AH, respectively.

This observation suggests that although our two-dimensional descriptor successfully identified membrane-permeabilizing candidates that are active on simplified model liposomes, only a small subset of these peptides will retain inhibitory activity against authentic viral envelopes because of other, external factors that were not incorporated in our present *in silico* and *in vitro* approaches. These insights highlight the need for advanced screening assays that more accurately capture *both* curvature-dependent binding *and* lipid compositional dependency [23], enabling discrimination between true AVPs and non-toxic analogues active only against simplified model membranes.

## Discussion

Selective viral membrane disruption is governed by a narrow thermodynamic design window defined by curvature affinity and hydrophobicity. In this regime—characterized by relative curvature-binding free energies (ΔΔ*F*) of approximately −10 to −25 kJ mol^−1^ and mean hydrophobicities ⟨ *H* ⟩ ≈ 0.5–1.0—amphipathic peptides generate sufficient membrane leaflet stress to permeabilize viral envelopes while avoiding excessive nonspecific binding and cytotoxicity. This thermodynamic framework rationalizes membrane-targeting antiviral peptides discovered empirically in earlier studies [16, 19]. The therapeutic value of sequences such as the hepatitis C virus amphipathic helix (HCV AH), identified serendipitously nearly two decades ago [16], illustrates how amphipathic helices exploit nanoscale membrane curvature to modulate biological function [22]. Related strategies have since extended beyond antiviral applications to cancer immunotherapy through the selective targeting of tumor-derived exosomes [44]. Despite these advances, general design principles governing curvature-dependent membrane activity have remained elusive. Purely data-driven AI approaches [26] are limited by the scarcity of experimentally validated curvature-sensitive sequences [22, 45], motivating physics-based strategies that directly encode interfacial thermodynamics

Over the last six years, we cycled through four generations of design approaches to systematically explore peptide sequence space. Rather than progressively optimizing antiviral potency, this iterative exploration enabled us to map how curvature affinity and hydrophobicity jointly determine membrane permeabilization and cytocompatibility. In total, we designed, synthesized, and tested 43 *de novo* candidates. Several displayed antiviral activity, with lead P1.6 achieving substantially higher potency against ZIKV than HCV AH in direct comparisons without showing toxicity, as characterized in more detail in our parallel work [23]. Most importantly, integrating newly generated and literature sequences allowed us to confine functional activity to a distinct thermodynamic regime characterized by strong membrane-permeabilizing potency while remaining non-cytotoxic.

To promote reproducibility and enable independent advancement of membrane-active peptide design, we projected peptide activity onto a thermodynamically intuitive two-dimensional descriptor space defined by affinity for positive membrane curvature and overall hydrophobicity (Fig. 4A). This mapping revealed a confined functional ‘sweet spot’ in which permeabilization, antiviral efficacy, and cytocompatibility intersect. Both descriptors can be computed in seconds for any user-provided sequence using our public PMIpred server (https://pmipred.fkt.physik.tu-dortmund.de) [25], ensuring full reproducibility and enabling community-driven navigation of this low-dimensional design space.

Viral selectivity likely exploits the absence of active membrane repair mechanisms in viruses, decoupling antiviral potency from liposome size–dependent permeabilization often observed in simplified model systems. Vero E6 cells derived from African green monkey kidney epithelium and TZM-bl cells – human engineered cervical epithelial cell – provide complementary mammalian models for evaluating peptide–membrane interactions [18, 20, 23]. Their cholesterol-rich membranes differ markedly from viral envelopes and constitute a stringent test for off-target cytolytic effects [46]. Peptides from our generation P4.x series, selected from within the identified ‘sweet spot’, exhibited robust activity against model membranes while remaining non-toxic to both TZM-bl and Vero E6 cells. These findings demonstrate that strong membrane permeabilization can be achieved without indiscriminate cytotoxicity when operating within an appropriate thermodynamic window.

At the same time, the contrast between high potency in simplified liposome assays and reduced efficacy against authentic viruses underscores the importance of membrane composition. Our detailed study of P1.6 demonstrated that its membrane-permeabilizing activity is markedly reduced by elevated levels of cholesterol or phosphatidylethanolamine (PE) [23]. The ternary model liposomes (e.g., DOPC:SM:cholesterol) used as viral mimics [32] tend to phase-separate into coexisting liquid-disordered and liquid-ordered domains [47], potentially concentrating peptides in cholesterol-depleted regions and thereby enhancing apparent permeabilization. In contrast, HIV-1 viral envelopes exhibit a uniform, cholesterol-enriched liquid-ordered phase [48]. Binary PC:cholesterol mixtures, which do not undergo such phase separation, more closely reproduce this uniform ordering and yield membrane-permeabilizing profiles consistent with observed antiviral efficacy [23]. These distinctions may explain why P1.6 outperforms HCV AH and C5A in several liposomal assays yet underperforms against HIV-1 [23], in line with atomistic simulations demonstrating that cholesterol enrichment markedly impairs its membrane binding ability (see Fig. 2G). Collectively, these observations indicate that accurate prediction of AVP activity requires membrane models that incorporate essential compositional features such as cholesterol-induced ordering and phosphatidylethanolamine enrichment [48, 49].

Building on these insights, we are developing an inverse lipidomics framework that leverages differential binding data from experimentally validated peptides—true antivirals (e.g., P1.6, HCV AH, C5A) versus liposome – only actives (e.g., P4.x) – to reconstruct target membrane compositions that best rationalize observed selectivity through differences in binding affinity [50]. This empirically guided strategy aims to refine membrane models used in computational optimization by correcting systematic model errors and simplifying assumptions. Integrating such composition-aware membrane representations with our curvature-based framework is expected to further enhance the predictive design of antiviral peptides while maintaining minimal host-cell toxicity.

Mechanistically, our results reinforce the thermodynamic link between curvature affinity and leaflet stress generation. The antiviral efficacy of active sequences such as P2.2.3 and P1.6 correlates with liposomal leakage activity and with their ability to stabilize highly positively curved toroidal pore structures in molecular simulations (Fig. SI14 and SI15). Transmission electron microscopy (TEM) further shows that P1.6 can remove the membrane envelope from viral capsids (Fig. 2C) [22, 23]. Our earlier work established the thermodynamic equivalence between leaflet stress generation and curvature affinity, implying that peptides optimized for positive curvature intrinsically encode membrane-permeabilizing propensity. Within the present framework, this propensity must be balanced against hydrophobicity to avoid excessive nonspecific membrane association.

Importantly, membrane-targeting antivirals need not rely exclusively on permeabilization. Amphiphilic peptides derived from influenza M2 [33] and claudin (CL) [34] inhibit viral infection by deforming envelopes without inducing lysis. These deformation-inducing sequences exhibit distinct curvature affinities and hydrophobicities relative to classical membrane-permeabilizing antivirals (Fig. 1A), suggesting that alternative functional regimes may exist outside the permeabilization sweet spot identified here. Such regimes may stabilize curvature stress without pore formation, further illustrating how low-dimensional thermodynamic descriptors may organize diverse membrane-active behaviors.

The continued development of physics-based generative design approaches therefore marks a conceptual shift from empirical sequence heuristics toward quantitative navigation of interfacial free-energy landscapes [39]. By coupling curvature-dependent binding with composition-aware membrane models, these frameworks provide a rational basis for predicting and directing peptide behavior across diverse viral envelope architectures. Selective membrane permeabilization and antiviral activity are confined to a narrow thermodynamic design window defined by curvature affinity and hydrophobicity, linking biological function to a low-dimensional free-energy landscape. More broadly, identifying a thermodynamic design window for membrane remodeling demonstrates how low-dimensional free-energy landscapes can be navigated to engineer selective interactions at soft-matter interfaces, providing a rational route toward membrane-active biomaterials and therapeutics.

## Data availability

All data supporting the findings of this study are available within the paper and its Supplementary Information. Additional simulation data and peptide sequence datasets are available from the corresponding authors upon reasonable request.

## Code availability

The PMIpred predictor used in this study is publicly accessible at https://pmipred.fkt.physik.tu-dortmund.de. Code used for evolutionary molecular dynamics simulations is available from the corresponding authors upon reasonable request.

## Methods

Detailed descriptions of the Evo-MD design framework, molecular dynamics simulations, free-energy calculations and experimental procedures are provided in the *Supplementary Methods* and *Supplementary Information*.

## Acknowledgements

The Dutch Research Organization NWO (Snellius@Surfsara) are acknowledged for provided computational resources. H.J.R. also gratefully acknowledges the Gauss Centre for Supercomputing e.V. (www.gauss-centre.eu) for funding this project by providing computing time through the John von Neumann Institute for Computing (NIC) on the GCS Supercomputer JUWELS at Jülich Supercomputing Centre (JSC) and the High-Performance Computing Center Stuttgart (HLRS). This work was supported by the German Research Foundation (DFG) through a grant to J.Mü. and H.J.R. (530287567), and to J.Mü. (CRC1279). N.v.H, J.Met. and H.J.R were also supported by the Dutch NWO Vidi scheme (project number 723.016.005). H.J.R additionally acknowledges the DFG under Germany’s Excellence Strategy-EXC 2033-390677874-RESOLV for support. MH acknowledges support by the the German Research Foundation (DFG 415894560) and the European Social Fund Plus (ESF+) together with the Federal State of Saxony-Anhalt (Grant No. ZS/2024/09/189789). Further funding to MH was contributed by Daimler and Benz foundation.

## Supplementary Methods (SM)

### 1 Methodology: Physics-Driven generative model

#### 1.1 Generation P1.x: The thermodynamic gradient approach

As shown previously for ALPS, curvature sensing by peptides can be studied by introducing an external thinning potential that locally induces lipid packing defects on a flat membrane [10]. Because the slope of the free energy in the packing defect gradient (“buffer zone”) is linear, the sorting propensity can be quantified by performing a single simulation with the peptide constrained in the middle of the defect gradient (Fig. SM1A), and directly measuring the sorting force 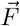, i.e. the spatial derivative of the potential of mean force, 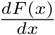 [10]. By definition, a negative sorting force indicates an attraction toward the defected thinned zone (i.e. ‘positive curvature’ sensing) and a positive sorting force would indicate a repulsion.

We implemented this force calculation as a fitness function in Evo-MD. To reduce the search space, the sequences in these runs were made up of recurring units of block length *L*_block_, that were repeated *r* times such that the total length (*r* × *L*_block_) varied from 2 to 24 amino acids (as further detailed in Ref. [23]). We observed a quick convergence of the average fitness to a sorting force of about -6.5 kJ mol^−1^ nm^−1^ (Fig. SM1B), with a clear preference for longer peptides since the sorting force is intrinsically proportional to peptide length.

To facilitate the intrinsic tendency for the longer peptides to dominate the population, we disabled the sequence-shortening operations after iteration 20, such that sequence lengths only increased from this point onward. Simultaneously, we limited the peptide net charge *z* to either slightly negative − 3 ≤ *z* ≤ 0, or slightly positive 0 ≤ *z* ≤ 3, or we left it unconstrained (red, blue, and black lines in Fig. SM1B, respectively). This allowed further improvement of the average fitness until the sorting force plateaued at about -9 kJ mol^−1^ nm^−1^ between iteration 20 and 40. Finally, to fine-tune the sequences in the population, Evo-MD was continued for another 10 iterations with disabled crossover operations, a double mutation rate and random residue swapping. For each of the three runs, the three most highly scoring peptides (named P1.x, all scoring <-11 kJ mol^−1^ nm^−1^) in the final population are described in Table SI4 and Fig. SI1.

Although we achieved evolutionary convergence, the thermodynamic gradient approach comes with two fundamental drawbacks. First, the sorting force strongly fluctuates over the MD trajectory due to the slow orientational and rotational modes of the peptide in the asymmetric system, which causes slow convergence and a relatively low signal-to-noise ratio. Moreover, since we shrunk the system size (compared to the setup in Ref. [10]) to reach the computational efficiency required for high-throughput evaluation in the Evo-MD framework, the lipid packing defect gradient is now within the same size range as the peptide itself (Fig. SM1A), which aggravates these effects. Second, the decision to reduce the search space by using repeated blocks of length *L*_block_ complicated the emergence of peptides with high amphipathicity, a feature that is well known to correlate with membrane surface adhesion [51], curvature sensing [52], and membrane disruption [33, 53]. This notion is illustrated by the selected hits in Fig. SI1: since all high-fitness sequences were 24 residues and the longest *L*_block_ was 12, all optimized peptides converged to a 12 × 2 structure. Because the number of AAs per helical turn is 3.6 and 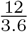 is not an integer, this always yields peptides that have a low overall hydrophobic moment.

**Fig. SM1.**
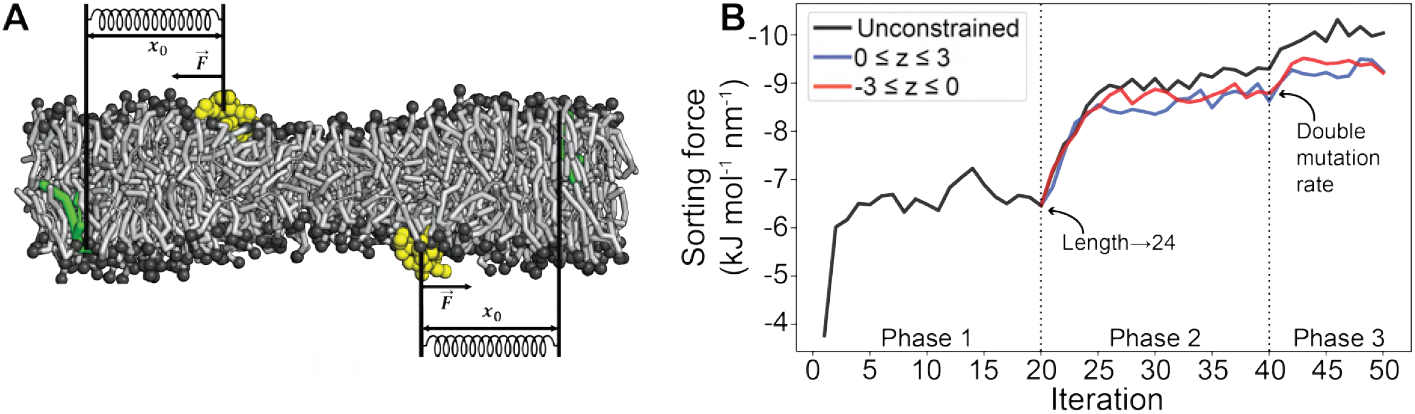
Sorting force-driven Evo-MD. **A)** System setup used for the Evo-MD runs with sorting force 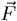 as the fitness function. POPC lipids in gray. Peptides in yellow. Reference POPC molecules in green. **B)** The average sorting force (fitness) of the population increases during three phases of simulated evolution. The dotted lines indicate the points at which the probabilities for the genetic operations are altered to facilitate fitness increase (see Ref. [23] for details). The *y*-axis is inverted to visually emphasize that we wish to maximize the magnitude of the sorting force. Figure adapted from Ref. [23].

In the next three peptide generations, we overcame both of these drawbacks: (1) we replace the sorting force-based fitness function with a more efficient and accurate end state free-energy calculation approach [24] (2) we only considered peptides with length 24, which allowed us to simplify the genetic operation scheme and only apply crossover recombination and point mutations.

#### 1.2 Generation P2.x: Optimizing sensing free energy while constraining the number of polar residues

Our previous work revealed that the best curvature-sorting peptides are equivalently the best curvature-generating peptides [9]. These optimal curvature-generators are extremely hydrophobic and bulky. Clearly, this physical optimum does not fulfill one of the main criteria of biological function: solubility in water. Therefore, we should narrow down our search space to the realm of biologically relevant water-soluble peptides.

A straight-forward way to increase the solubility of a peptide is to introduce polar residues (D, E, H, K, N, Q, R, S, or T). We performed three runs of Evo-MD with the same settings as in Ref. [9], whilst – one-the-fly – discarding any sequences with fewer than 8 polar residues that arose during evolution. All three runs converged to optima with |ΔΔ*F*| values of around 26 kJ mol^−1^, 6 kJ mol^−1^ lower than the global optima without any polar constraints (Fig. SM2A). Naturally, this is due to bulky hydrophobic residues (F/W) being replaced by polar ones (mainly K/E/Q), which reduces the peptide’s ability to maximally induce leaflet tension upon binding. When we continued the Evo-MD run with even more stringent constraints (having at least 10 or 12 polar residues), the |ΔΔ*F*| of the optima drops to lower values, in line with this trend (Fig. SM2A).

We note that enforcing *at least n* polar residues lead to solutions with *exactly n* polar residues. In other words, the number of polar residues is always minimized, as was the case in the unconstrained runs. As evident from the helical wheel representations of the second generation of peptides (named P2.x, see Fig. SM2B-E and Fig. SI2), the newly introduced polar AAs tend to cluster at the same helical face, hereby increasing the peptide’s amphipathicity (hydrophobic moment *µ*_H_) with respect to the unconstrained optima.

**Fig. SM2.**
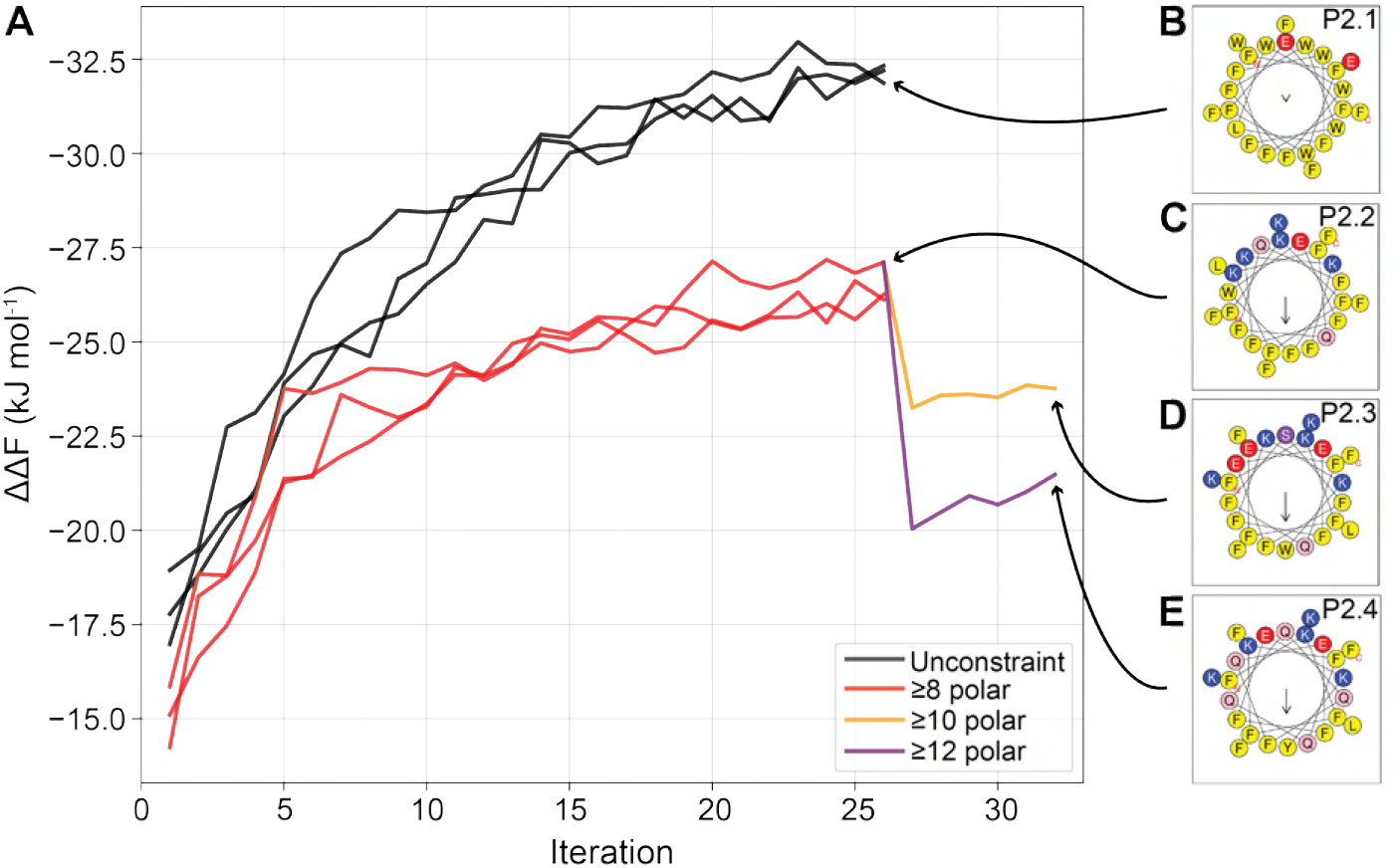
Evo-MD runs with polar constraints. **A)** Evo-MD convergence of the best candidates for every population without constraining the number of polar residues (in black), with at least 8 polar residues (in red), at least 10 polar residues (in orange), or at least 12 polar residues (in purple). The *y*-axis is inverted to visually emphasize that we wish to maximize the magnitude of ΔΔ*F*. **B-E)** Helical wheel representations [37] of the resolved optima P2.1, P2.2, P2.3, and P2.4 (see Table SI1 and Fig SI2 for details). Yellow: hydrophobic. Pink/purple: polar. Blue: positively charged. Red: negatively charged.

#### 1.3 Generation P3.x: Multi-objective optimization

By introducing strict rules on the minimal number of polar residues in the sequences, we managed to design peptides that were less hydrophobic and more amphipathic than the absolute optima. However, mean hydrophobicities are still very high for these peptides (⟨ *H* ⟩ >0.75 for most sequences, see Table SI1). Higher hydrophobicity naturally means poorer solubility, which may complicate peptide synthesis and experimental characterization. Moreover, even when such peptides are sufficiently soluble, they may not interact with membranes at all, as we have seen for peptide candidate P2.2 (Fig. SI13B), likely due to premature self-aggregation. To overcome this hurdle, we introduced the minimization of mean hydrophobicity as a second objective in the optimization process, much like natural evolution of proteins would.

In a multi-objective optimization, one defines *N* objectives that are desired to be optimized simultaneously. Rather than the optimum being a single value (as we have seen in the previous Evo-MD optimizations), the optimum is now defined as an (*N*-1)-dimensional ‘pareto-front’: i.e. a set of candidate solutions that are considered equally optimal. Here, we wish to maximize |ΔΔ*F*| and minimize the mean hydrophobicity ⟨ *H* ⟩. Thus, we are dealing with a dual-objective optimization problem (*N* = 2) and, therefore, the pareto front is a curve, which’ extrema represent the optimal values for the individual objectives. All solutions between these extrema comprise a unique trade-off between the two objectives.

We implemented this multi-objective approach in Evo-MD using the NSGA-II algorithm [31] to rank the candidate solutions in a population after every iteration of the algorithm. Additionally, to narrow down the search space to the realistic realm of membrane-surface-active, yet water-soluble peptides, we used the antimicrobial peptide database (APD) [54] to define peptide property ranges that all generated sequences should obey (e.g. rules for AA content, net charge, and hydrophobic moment, see Fig. SM3A). Finally, we considered all 20 AAs within the evolution of peptides, but taking into account their respective helical propensities [55] during the initial sequence generation and all point mutation steps (Table SM1). All other settings were the same as in previous single-objective runs conducted in Ref. [9].

When we ran this dual-objective Evo-MD, we observed that the pareto front indeed converged to the upper left corner of the 2D-solution space where it has a high |ΔΔ*F*| and a low ⟨ *H* ⟩ (Fig. SM3B), as intended. Evolutionary convergence plots of the separate objectives show that ΔΔ*F* was relatively stable over the course of evolution, with the best solutions at about -20 kJ mol^−1^ (Fig. SM3C). Interestingly, it was mostly the mean hydrophobicity ⟨ *H* ⟩ that decreased (for a constant ΔΔ*F*) until evolution converged to an optimal value of -0.4 (Fig. SM3D).

The pareto front of the final iteration (Fig. SI3A) comprises peptides that span rather wide ranges of ΔΔ*F* (roughly -5 to -19 kJ mol^−1^) and ⟨ *H* ⟩ (roughly -0.4 to 0.5). Sequences, properties, and helical wheel plots of these peptides, named P3.x, are provided in Table SI1 and Fig. SI3B. Within our thermodynamic membrane-binding model, these peptides are spread out across the lower binding regime that thus far yielded the most promising results (e.g., P1.6, P2.2.3, and positive control HCV AH), except P3.5 d and P3.6 which are predicted to be close to the non-binder → sensor boundary (Fig. SI4).

**Fig. SM3.**
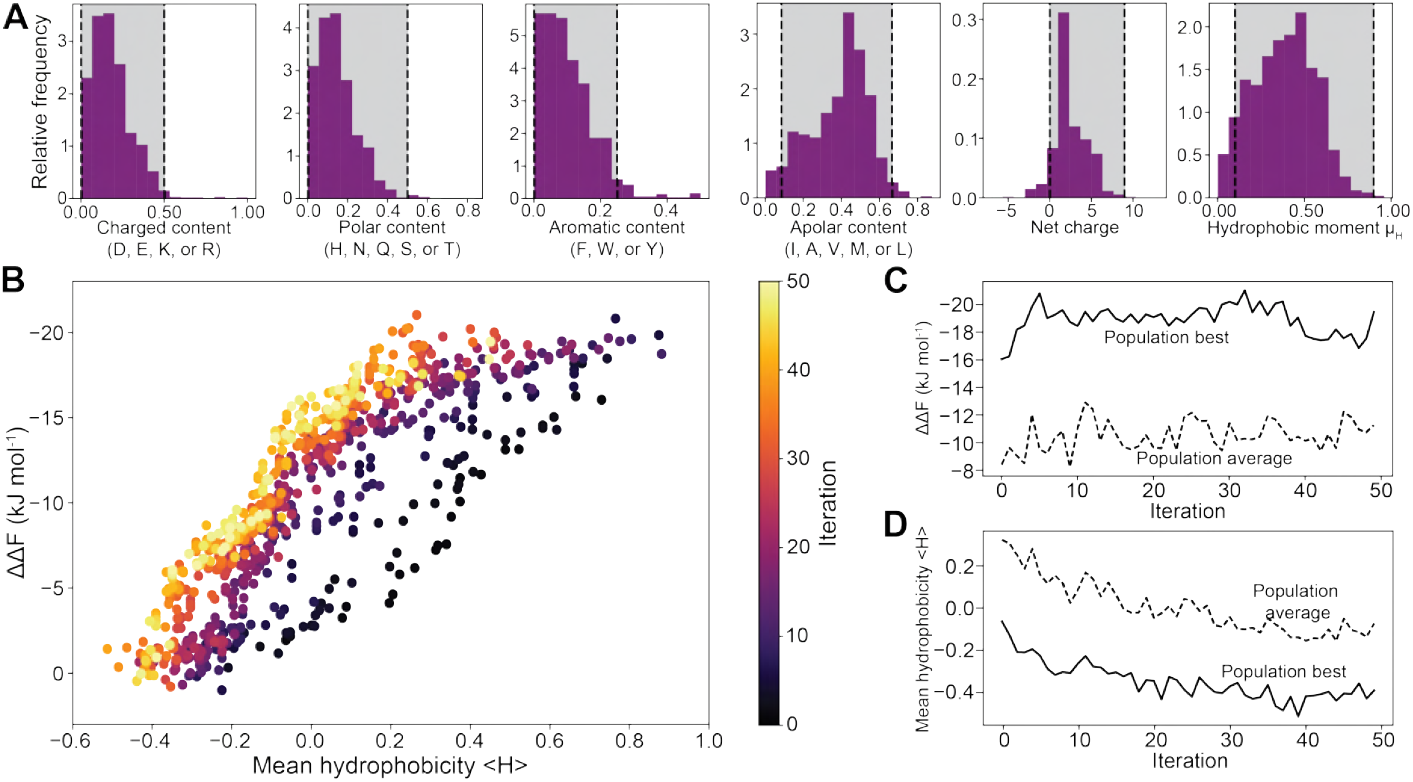
Dual-objective Evo-MD. **A)** Histograms showing the distributions of charged content (relative number of D, E, K, or R residues), polar content (relative number of H, N, Q, S, or T residues), aromatic content (relative number of F, W, or Y residues), apolar content (relative number of I, A, V, M, or L residues), net charge, and hydrophobic moment for all 1,256 sequences of length ≤24 in APD3[54]. The gray area and black dotted lines indicate the regime in which Evo-MD-generated sequences should fall in order to be accepted by the algorithm. **B)** Scatter plot of the first pareto front *P*_1_ of every Evo-MD iteration for the two objectives ΔΔ*F* and ⟨ *H* ⟩. Brighter colors indicate later iterations. **C)** ΔΔ*F* progression over 50 iterations of dual-objective Evo-MD. **D)** Mean hydrophobicity ⟨ *H* ⟩ progression over 50 iterations of dual-objective Evo-MD.

**Table SM1:** *α*-helix propensities. Relative probabilities for the 20 natural AAs derived from the experimentally determined helical propensities (relative to Ala) [55] by the Boltzmann distribution 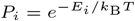.

| AA | Helix propensity (kJ mol<sup>-1</sup>) | Probability |
| --- | --- | --- |
| Ala (A) | 0.00 | 0.110 |
| Leu (L) | 0.88 | 0.077 |
| Arg (R) | 0.88 | 0.077 |
| Met (M) | 1.00 | 0.073 |
| Lys (K) | 1.09 | 0.071 |
| Gln (Q) | 1.63 | 0.057 |
| Glu (E) | 1.67 | 0.056 |
| Ile (I) | 1.72 | 0.055 |
| Trp (W) | 2.05 | 0.048 |
| Ser (S) | 2.09 | 0.047 |
| Tyr (Y) | 2.22 | 0.045 |
| Phe (F) | 2.26 | 0.044 |
| Val (V) | 2.55 | 0.039 |
| His (H) | 2.55 | 0.039 |
| Asn (N) | 2.72 | 0.037 |
| Thr (T) | 2.76 | 0.036 |
| Cys (C) | 2.85 | 0.035 |
| Asp (D) | 2.89 | 0.034 |
| Gly (G) | 4.18 | 0.020 |
| Pro (P) | 13.22 | 0.001 |

#### 1.4 Generation P4.x: Sweet spot approach

The fourth generation of peptides was obtained by reassessing a large number of sequences generated in Generations 2 and 3 on their respective coordinates in ⟨ *H* ⟩, ΔΔ*F*_*adj*_ space. First, we identified the sweet spot region where we anticipated active antiviral peptides would be located on the basis of the previous 3 generations as well as the known examples from literature. A systematic analysis was conducted to identify peptides within this predefined elliptical region (Fig SM4) using the generated database consisting of 53,940 sequences [9]. These generated data points strategically cover the whole two-dimensional descriptor space {⟨*H*⟩, ΔΔ*F*}.

Twenty peptide sequence were selected guided by two primary criteria:

- Proximity requirement: Euclidean distance less than 0.1 units from a known active sequence.
- Energy parameter alignment: Difference in PMIpred-predicted ΔΔ*F* and MD ΔΔ*F* values below 1.0 kJ/mol

From the qualifying peptides at each point, selection was based on two key charge-based parameters that have proven successful in generation P1.x and P2.x:

- Highest charge moment (*µ*_*z*_) value
- Neutral or positive charge

An additional peptide was identified near the P1.6 and HCV AH regions, characterized by a negative charge (-1) and designated as P4.9. Ultimately, 10 of these peptides were selected for experimental validation.

**Fig. SM4.**
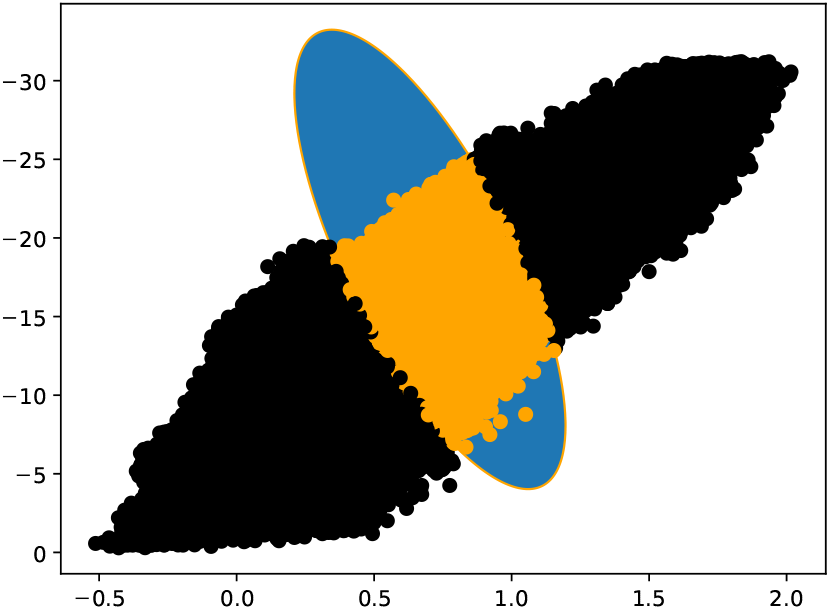
Sweet spot selection approach. Every data point represents one of the 53,940 sequences in {⟨ *H* ⟩ }, ΔΔ*F* space. The sweet spot region is marked by the blue ellipse. Sequences that fall within this region are marked orange.

### 2 General molecular dynamics settings

#### 2.1 Evolutionary Molecular Dynamics (Evo–MD)

Peptide sequences were generated using an evolutionary molecular dynamics (Evo–MD) framework as described previously [9, 24]. Evo–MD combines coarse-grained molecular dynamics simulations with a genetic optimization algorithm to evolve peptide sequences toward a predefined fitness function. In each iteration, peptide candidates were evaluated in a lipid membrane environment and ranked according to their relative curvature affinity ΔΔ*F*, defined as the difference in membrane binding free energy between a planar tensionless membrane and a membrane enriched in lipid packing defects mimicking positive curvature.

The first peptide generation (P1.x) was simulated using the Martini 2 force field. All subsequent peptide generations (P2.x, P3.x, and P4.x) were simulated using Martini 3. All simulations were performed with GROMACS [56]. POPC bilayers were used as model membranes. Curvature effects were introduced either by applying membrane thinning protocols or lateral strain to enhance lipid packing defects, following the methodology described in Ref. [9].

Simulations were conducted under NPT conditions at 310 K using a velocity-rescale thermostat with *τ*_*T*_ = 1 ps and semi-isotropic pressure coupling using a Berendsen barostat with *τ*_*P*_ = 4 ps and compressibility 4.5 × 10^−5^ bar^−1^. Unless otherwise stated, production runs employed a 20 fs time step. For restrained equilibration or enhanced sampling stages, a reduced time step of 10 fs was used. Flat-bottom restraints in the *z*-direction were applied where necessary to control peptide positioning relative to the membrane.

Relative binding free energies ΔΔ*F* reported in Table 1 were obtained using the PMIpred framework [25], which was trained on large-scale simulation data generated using a mechanical end-state free energy approach [24]. The values of Δ*F*_∞_ explicitly denoted as “simulation”, they were computed independently using Thermodynamic Integration (TI). The TI protocol is described below.

#### 2.2 Calculation of Δ*F*_∞_ using Thermodynamic Integration (TI)

##### System preparation

Thermodynamic integration calculations for the membrane-bound state were performed in a tension-less POPC bilayer system (6 × nm 6 × nm 10 × nm) containing 128 lipids (64 per leaflet) and approximately 1700 water beads. As a reference for the solvation free energy, a pure water box (7 nm × 7 nm × 10 nm) containing approximately 3100 water beads was constructed. All TI simulations were performed using the Martini 3 force field. Counterions were added to ensure charge neutrality.

##### Equilibration

Systems were energy-minimized and equilibrated under NPT conditions at 310 K using the same thermostat and barostat settings described above. Flat-bottom restraints in the *z*-direction were applied to maintain the peptide position relative to the membrane. A final equilibration was performed using Langevin dynamics for 10 ns with a 10 fs time step.

##### Integration procedure

Binding free energies (Δ*F*_∞_) were computed by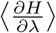 integrating with respect to *λ*. Van der Waals and Coulomb interactions were decoupled sequentially using 37 *λ* windows each (74 replicas in total). Production simulations of 500 ns were performed per *λ* value.

The free energy difference was calculated using trapezoidal integration:

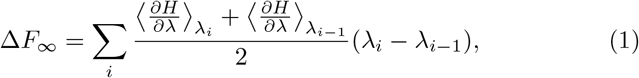

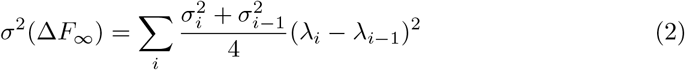

The results of the first three generations that included all highly active antiviral peptides are plotted in figure SM5.

**Fig. SM5.**
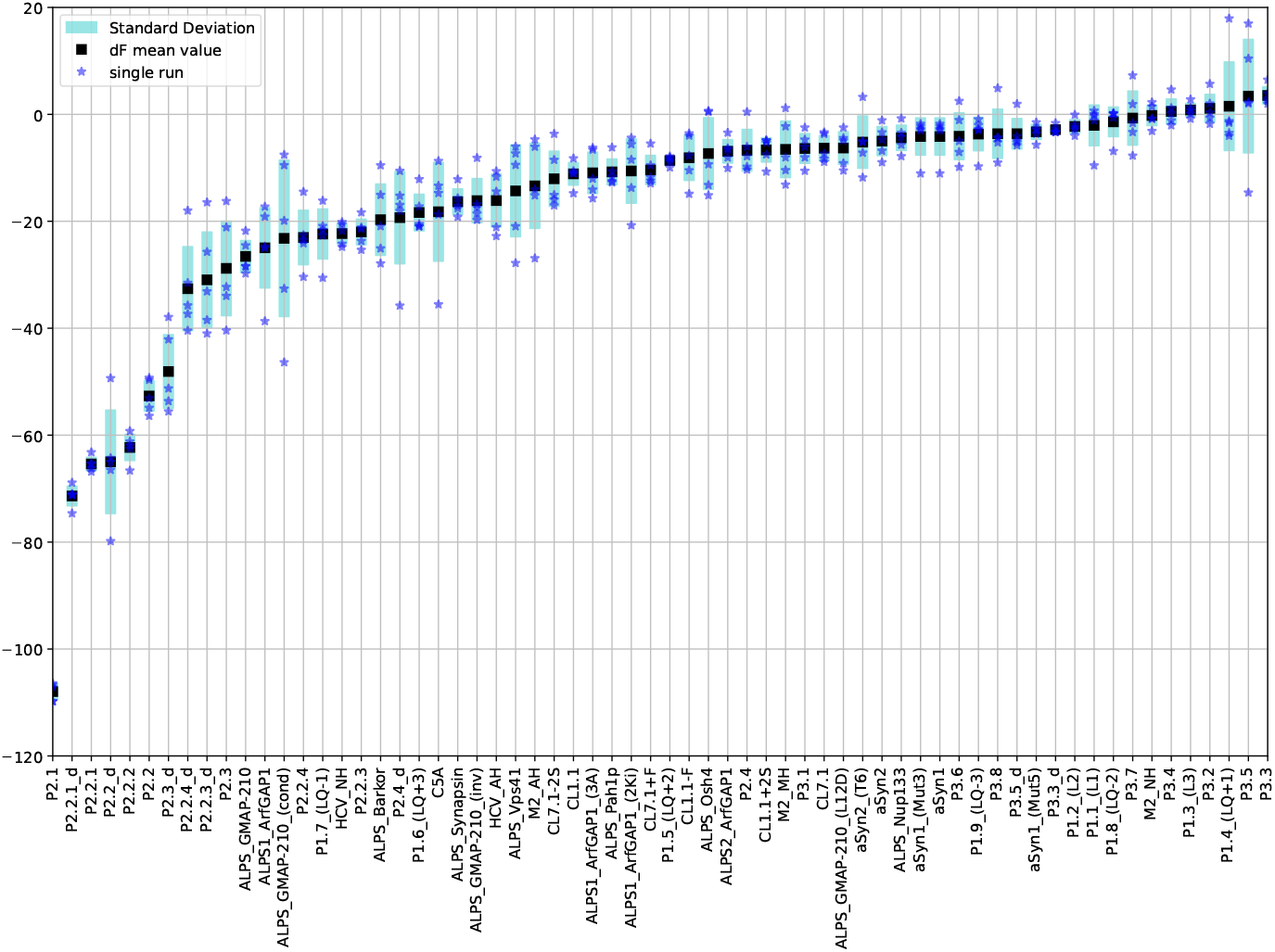
Explicit calculation of. Δ*F*_∞_. Shown are the Δ*F*_∞_ values calculated for peptide binding to a tensionless POPC membrane. The peptides from the first three generations are ordered from strongest membrane binding (large negative values) to weakest binding (values approaching zero or slightly positive).

### 3 Experimental methods and materials

#### Peptide synthesis

All peptides P1.x and P2.x were synthesized by solid-phase peptide synthesis using standard Fmoc-chemistry protocols. The synthesis was carried out on an automated Liberty Blue microwave peptide synthesizer (CEM). For deprotection of the Fmoc group 20% piperidine in dimethylformamide (DMF) was used, N,N’-diisopropylcarbodiimide (DIC)/Oxyma Pure were applied as the activator/base. All sequences were synthesized on a Tentagel R RAM resin. The coupling of AAs was performed at 90 °C for 4 min. The terminal amine was acetylated by shaking the resin with acetic anhydride (5 eq.) and pyridine (5 eq.) in DMF for 1 h. Afterwards the resin was washed 3 times with dichloromethane (DCM) and DMF. Subsequently the peptides were cleaved from the resin, using trifluoroacetic acid (TFA) with 2.5% deionized water and 2.5% triisopropylsilane (TIS). The crude peptides were precipitated in ice cold diethyl ether (Et_2_O), pelleted by centrifugation and blow dried with nitrogen. The crude was then redissolved in water and lyophilized. The peptide purification was performed on a Shimadzu HPLC system equipped with two LC-20AR pumps, an SPD-20A UV-Vis detector. For purification a Phenomenex Kinetex EVO C18 column (21.2 mm by 150 mm) was used. The mobile phases were water (H_2_O) and acetonitrile (ACN), containing 0.1% TFA. All purified peptides were lyophilized and stored at -20 °C until use.

All peptides P3.x and P4.x were purchased from the Core Facility Functional Peptidomics at the Ulm University Medical Centre, with the N-terminus acetylated and the C-terminus aminated, following a similar solid-phase peptide synthesis protocol.

##### ZIKV assay

6,000 Vero E6 cells were seeded on day 1. Peptides were diluted to 200 *µ*M and titrated 1:3 in a u-well plate. Next, the peptides were added 1:1 (v/v) to a ZIKV dilution and incubated for 30 min at 37 °C. The virus/peptide mixture was added to the cells in a 1:5 dilution, after which the cells were incubated for 2 days at 37 °C. Analyses were performed with the ZIKV in-cell ELISA assay, as described by Müller *et al* [57]. All experiments were performed in technical triplicates (*R* = 3) and repeated in 1–3 independent experiments (*N* = 1 − 3), as indicated in the respective figure captions.

##### HIV-1 assay

10,000 TZM-bl reporter cells were seeded on day 1. Peptides were diluted to 200 *µ*M and titrated 1:3 in a u-well plate. Next, the peptides were added 1:1 (v/v) to a HIV-1 dilution and incubated for 30 min at 37 °C. The virus/peptide mixture was added to the cells in a 1:5 dilution. Infected cells were incubated for 3 days at 37 °C, and then measured via a *β*-Gal assay (ThermoFisher catalog number T1027) according to the manufacturer’s instructions. All experiments were performed in technical triplicates (*R* = 3) and repeated in 1–3 independent experiments (*N* = 1 − 3), as indicated in the respective figure captions.

##### Cell viability assay

Cells were seeded on day 1 with cell types matching the corresponding ZIKV (Vero E6) and HIV-1 (TZM-bl) assays. In line with the virus assays, peptides were diluted to 200 *µ*M and titrated in a u-well plate. Next, the peptides were added 1:1 (v/v) to pure DMEM cell media and incubated for 30 min at 37 °C, after which this mixture was added 1:5 to the cells. After incubating for 2/3 days (matching the corresponding virus assay) at 37 °C, the cells were measured via a CellTiter-Glo assay (Promega catalog number G7570) according to the manufacturer’s instructions. Each measurement was performed with three technical replicates (*R* = 3) in 1–2 independent experiments (*N* = 1–2), as indicated in the respective figure captions.

##### Hemolysis assay

5 mL of fresh blood was collected and centrifuged at 1,000 g for 10 min. Blood serum was removed and the number of erythrocytes/mL was determined. A dilution series was made and measured on the plate reader (at 405 nm) after addition of PBS and Triton X-100 to determine the signal-to-noise ratio. Next, erythrocytes were diluted to 1.44 *·* 10^8^ cells in 40 *µ*L in a V-well plate. Peptides were diluted to 1 mM, titrated 1:2 in a u-well plate, and then added 1:5 (v/v) to the erythrocytes, such that the highest concentration is 200 *µ*M. The solution was incubated while shaking at 500 rpm for 30 min at room temperature. Then, the plate was centrifuged at 1,500 rpm for 5 min. The supernatant was carefully pipeted into a fresh flat-well plate and measured on the plate reader at 405 nm. For all peptides, two replicate measurements (*R* = 2) were performed in one experiment.

##### Liposome dye leakage assay

Model liposomes (45/25/30 mol% DOPC /SM/cholesterol) were prepared with encapsulated fluorescein dye as described by Weil *et al* [32]. Fluorescence was examined with the Synergy fluorescence spectrometer for 5 minutes to determine the baseline fluorescence. Peptides were diluted to 30 *µ*M, titrated 1:3, added 1:10 to the liposomes, and then incubated for 30 min at 37°C. The fluorescence was recorded continuously for 30 min on the Synergy fluorescence spectrometer, after which 0.1% Triton X-100 was added to the solution to determine the maximal leakage (measured for 5 min). For all peptides, three replicate measurements (*R* = 3) were performed in one experiment.

##### Monolayer adsorption assay

The lipid 1,2-dimyristoyl-*sn*-glycero-3-phosphocholine (DMPC) was purchased as lyophilized powder or in chloroform from Avanti Polar Lipids (Alabaster, AL, USA) and used without further purification. NaCl, NaOH, and HCl (all pro analysis grade), chloroform, and methanol (both HPCL grade) were obtained from Carl Roth GmbH (Karlsruhe, Germany). All solutions were prepared using ultrapure water with a resistivity above 18.2 MΩ cm.

Interactions of peptides with lipid monolayers were examined with circular polytetrafluoroethylene (PTFE) Langmuir troughs equipped with Dyne Probe sensors (Kibron Inc., Helsinki, Finland) with a Kibron Delta Pi 4 (Kibron Inc., Helsinki, Finland). The aqueous subphase contained 150 mM NaCl at a pH of 7.4 and was thermostated at 25.0 °C (Julabo GmbH, Seelbach, Germany) and the setup was contained in a box. The troughs were soaked for 15 min with aqueous Hellmanex solution (5% v/v) (Carl Roth GmbH, Karlsruhe, Germany) and extensively rinsed (ultrapure water, Milli-Q Advantage A10, resistivity 18.2 MΩ cm, Merck Millipore, Darmstadt, Germany). The Dyne probe sensors were calibrated for each recording. DMPC was spread from a chloroformic solution (approximately 1 mM) onto the subphase. After evaporation of the solvent, the lipid monolayer was allowed to equilibrate for 20 min. The peptide solution was injected into the subphase through lateral injection holes. The final peptide concentration in the subphase was 0.2 *µ*M, 2 *µ*M or 5 *µ*M. The surface pressure was recorded over time. The maximal change in surface pressure Δ*π* was determined in equilibrium after approximately 600 to 800 min, or 200 min for cases with no apparent insertion, such as P1.6 interacting with the buffer-air interface or P.2.2, respectively. Origin 16 (Origin Lab Corp., Northampton, USA) was used for data analysis.

## Supplementary Information (SI)

### 1 Overview tables for all four peptide generations

**Table SI1:**
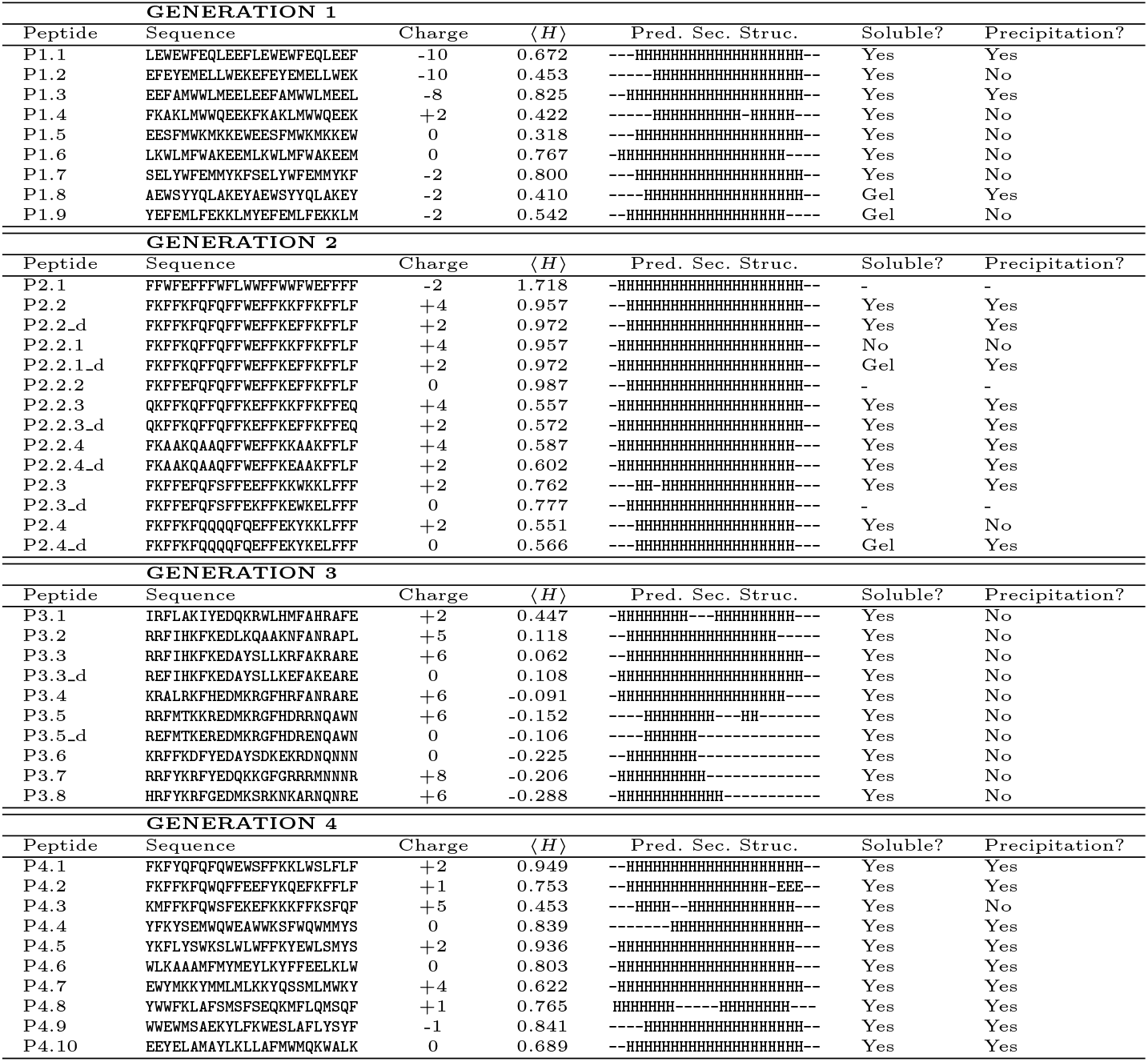
Peptide sequences. The P1.x sequences result from a sorting force-driven Evo-MD (section SM1.1). P2.1, P2.2, P2.3, and P2.4, are the best sequences from Evo-MD optimization runs with at least 0, 8, 10, and 12 polar residues, respectively (section SM1.2). All other P2.x sequences are derivatives of those four peptides. P3.x sequences were selected from the first Pareto front of dual-objective optimization Evo-MD (section SM1.3, Fig. SI3). P4.x peptides were selected to populate the here-defined AVP ‘sweet spot’ (section SM1.4, Fig. 4A) .The label ‘ d’ indicates mutations to avoid twin K motifs that can be involved in undesired targeting [58]. Helical wheel representations for all peptides are shown in Fig. SI1, SI2, SI3B, SI5). ⟨ *H* ⟩ : mean hydrophobicity; Pred. Sec. Struc.: predicted secondary structure, per JPred4 [59]; Solubility in DMSO/DMF; Precipitation in PBS.

**Table SI2:**
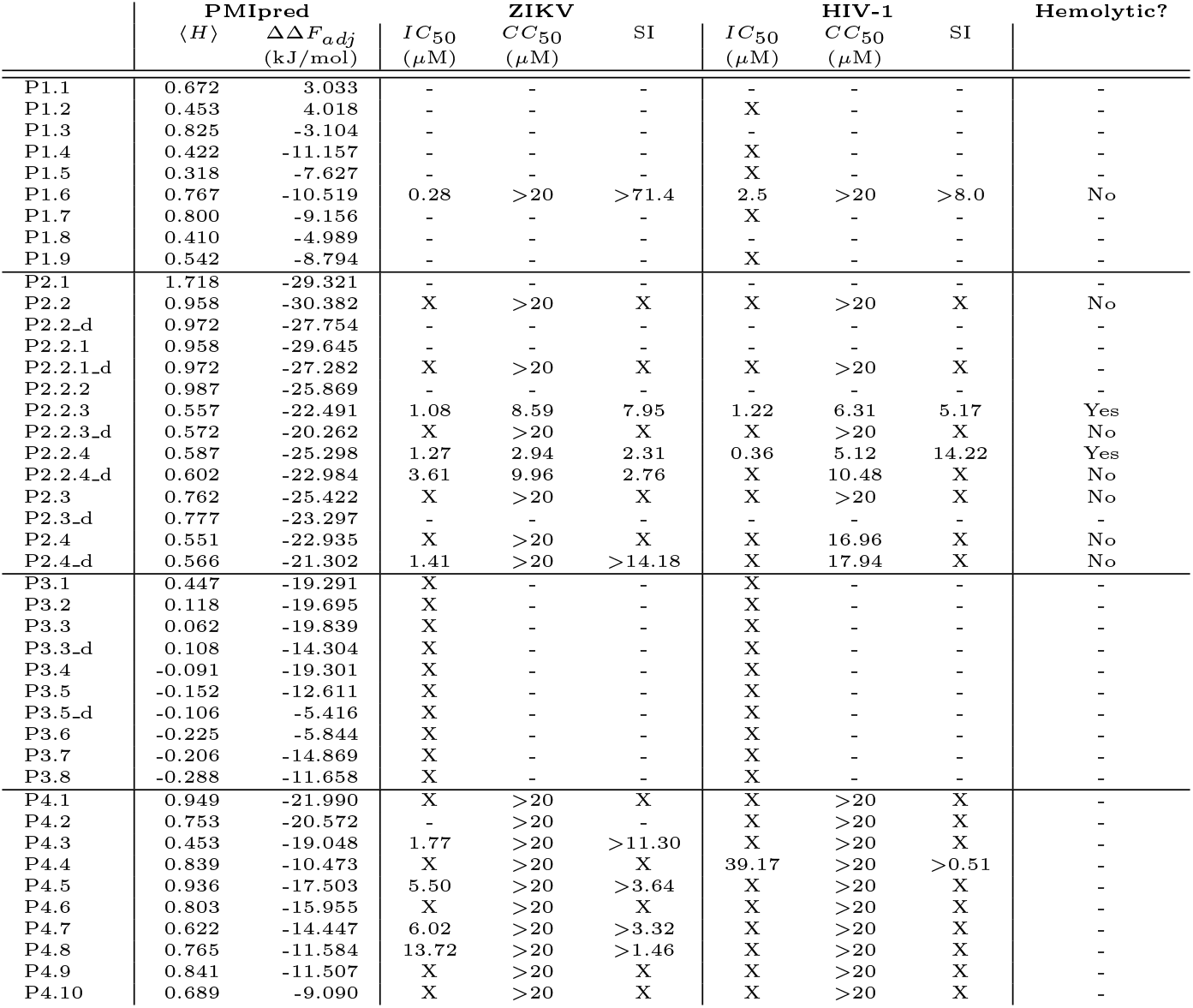
Overview of antiviral properties. Summary of data in Fig. 2A,H, Fig. SI6, Fig. SI7, Fig. SI8, Fig. SI9, Fig. SI10, Fig. SI11, Fig. SI16, Fig. SI17, Fig. SI18, Fig. SI19, and Fig. SI20. Data for P1.6 are taken from Ref. [23]. SI: selectivity index, i.e. the ratio *CC*_50_*/IC*_50_. X = inactive. - = not tested.

**Table SI3:**
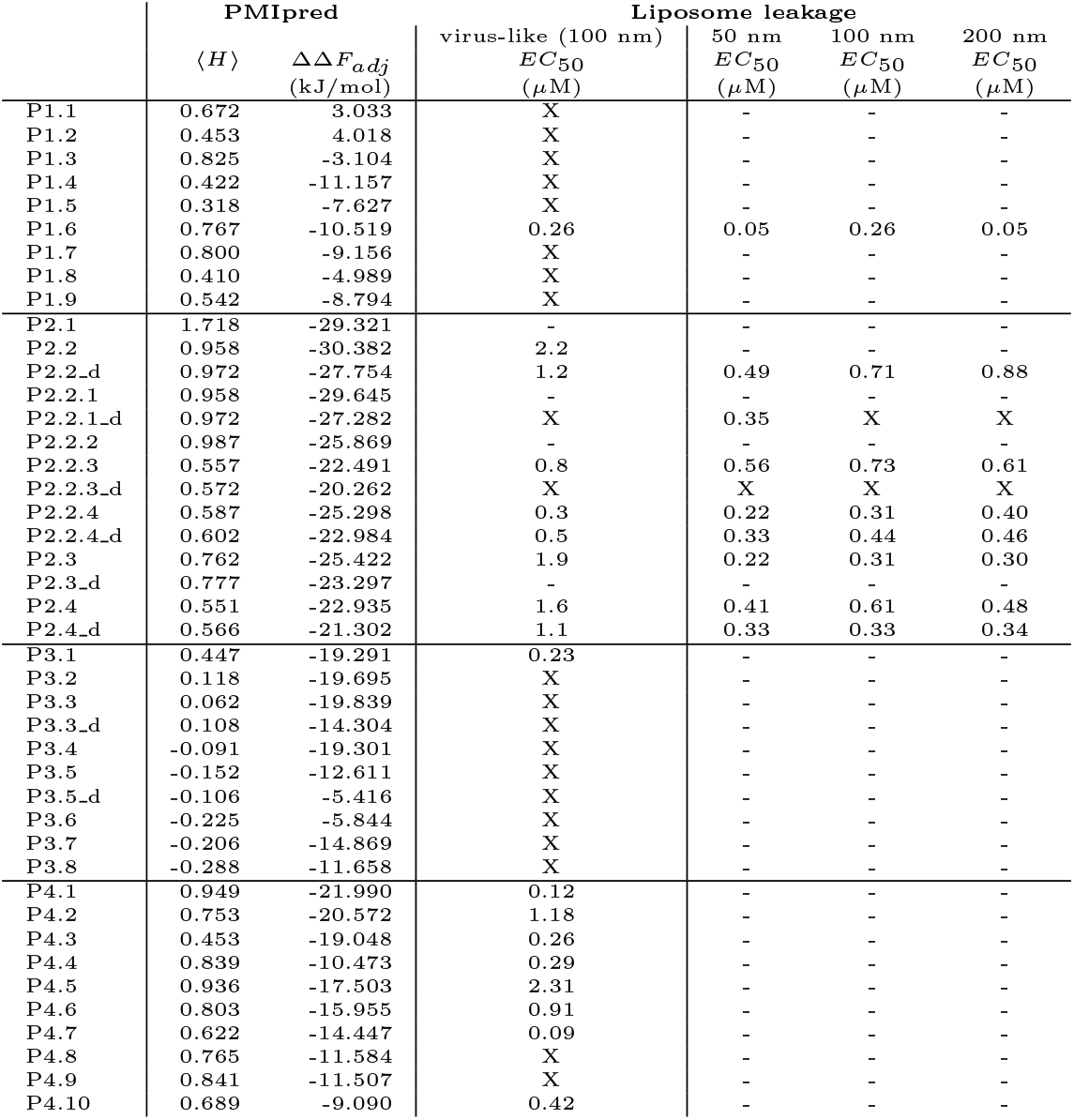
Overview of liposome leakage assay results. Summary of data in Fig. 2D-F, Fig. SI12. All liposomes contain 45/25/30 mol% DOPC/S-M/cholesterol, as described in Ref. [32]. Data for generation P1.x are taken from Ref. [23]. Virus-like liposomes are identical to 100 nm-sized liposomes. The EC_50_ values can differ for 100 nm liposomes (comparing liposome sizes) and virus-like liposomes because these experiments were performed independently. X = inactive. - = not tested.

### 2 Peptide selection

**Table SI4:**
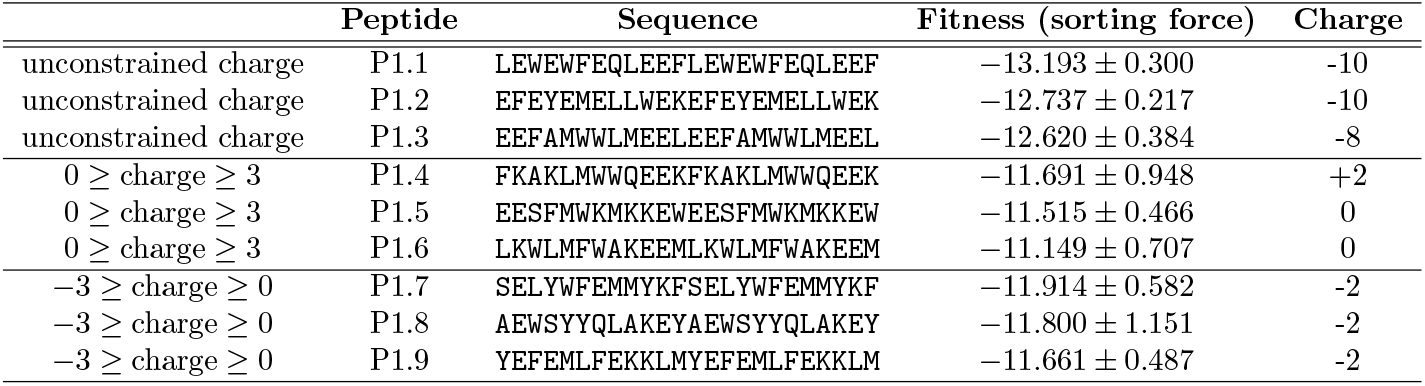
Fitness and peptide properties (generation 1) Sorting force fitness (kJ mol^−1^ nm^−1^) and net charge values for the selected peptides from the final populations of the sorting force-driven Evo-MD runs (Fig. SM1B). Helical wheel representations for all peptides are shown in Fig. SI1.

**Fig. SI1.**
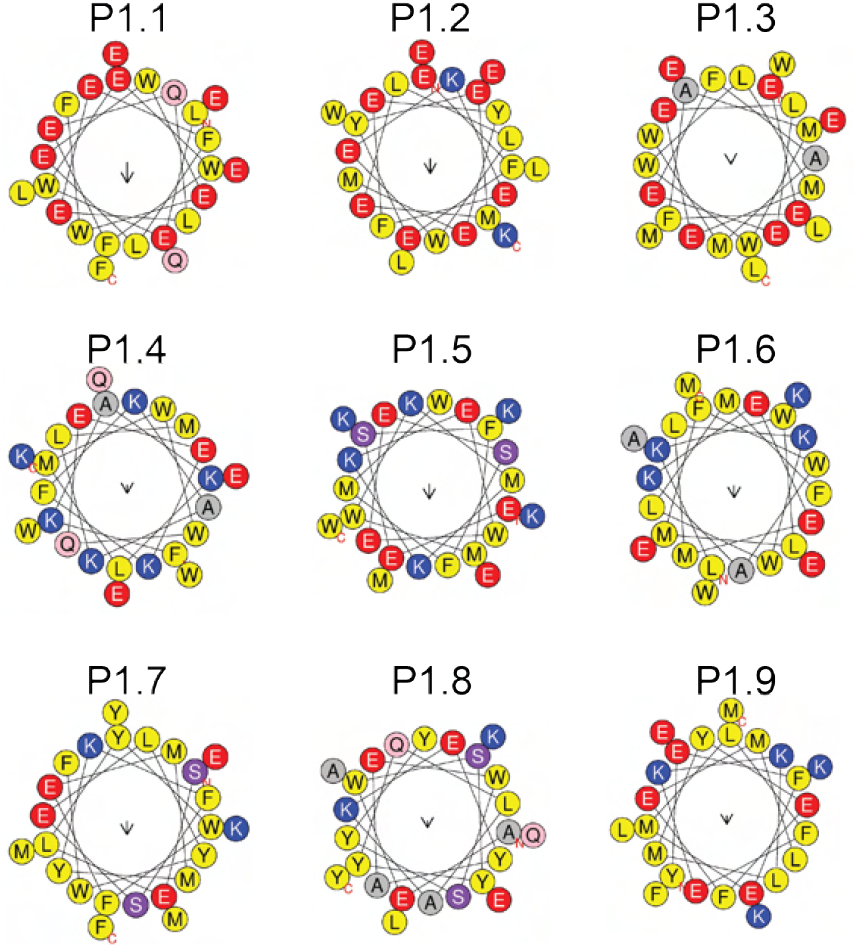
Helical wheels (generation 1). Helical wheel representations [37] for all P1.x peptides. Yellow: hydrophobic. Gray: small. Pink/purple: polar. Blue: positively charged. Red: negatively charged.

**Fig. SI2.**
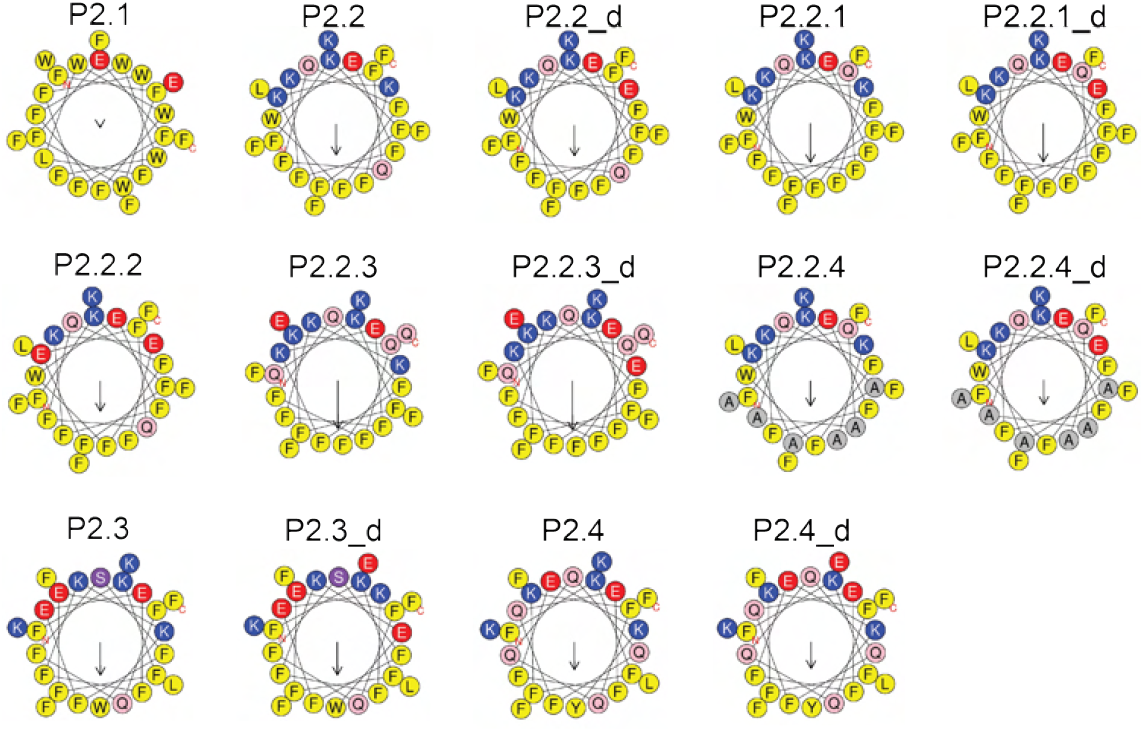
Helical wheels (generation 2). Helical wheel representations [37] for all P2.x peptides. Yellow: hydrophobic. Gray: small. Pink/purple: polar. Blue: positively charged. Red: negatively charged.

**Fig. SI3.**
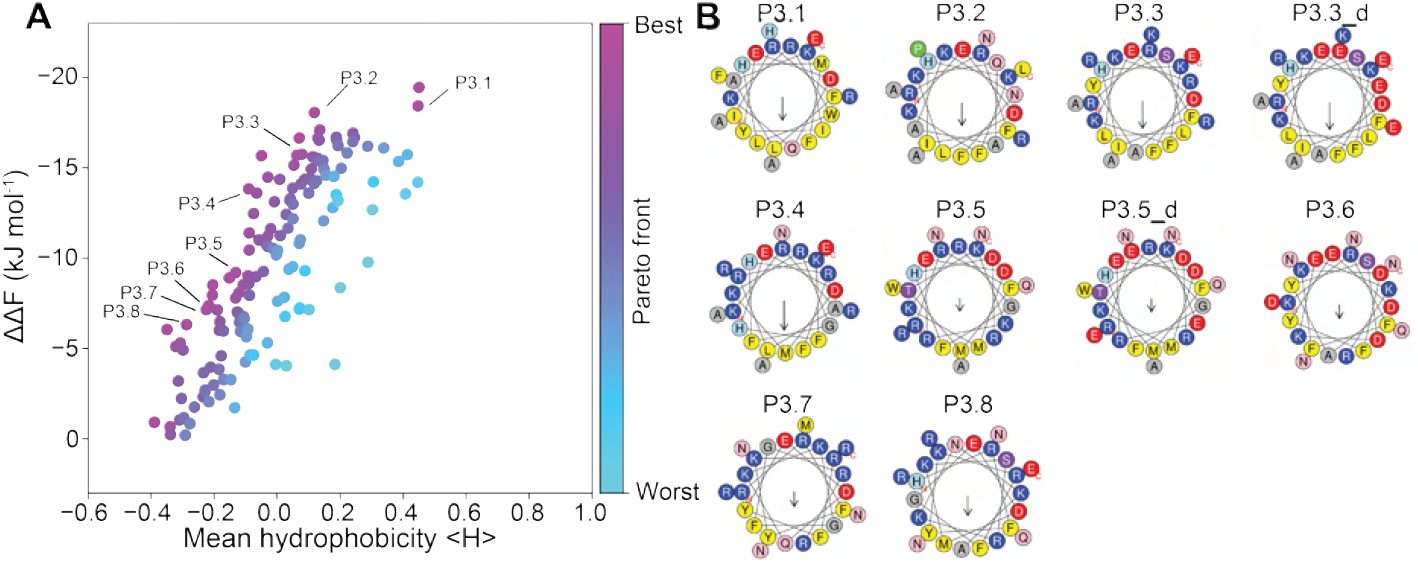
Details on the final iteration of multi-objective Evo-MD. **A)** The final population of the multi-objective Evo-MD run in Fig. SM3B. Pareto fronts *P*_*n*_ are colored from worst to best in cyan to magenta, respectively. **B)** Helical wheel representations [37] for all P3.x peptides. Yellow: hydrophobic. Gray: small. Pink/purple/loght blue: polar. Blue: positively charged. Red: negatively charged. Green: proline.

**Fig. SI4.**
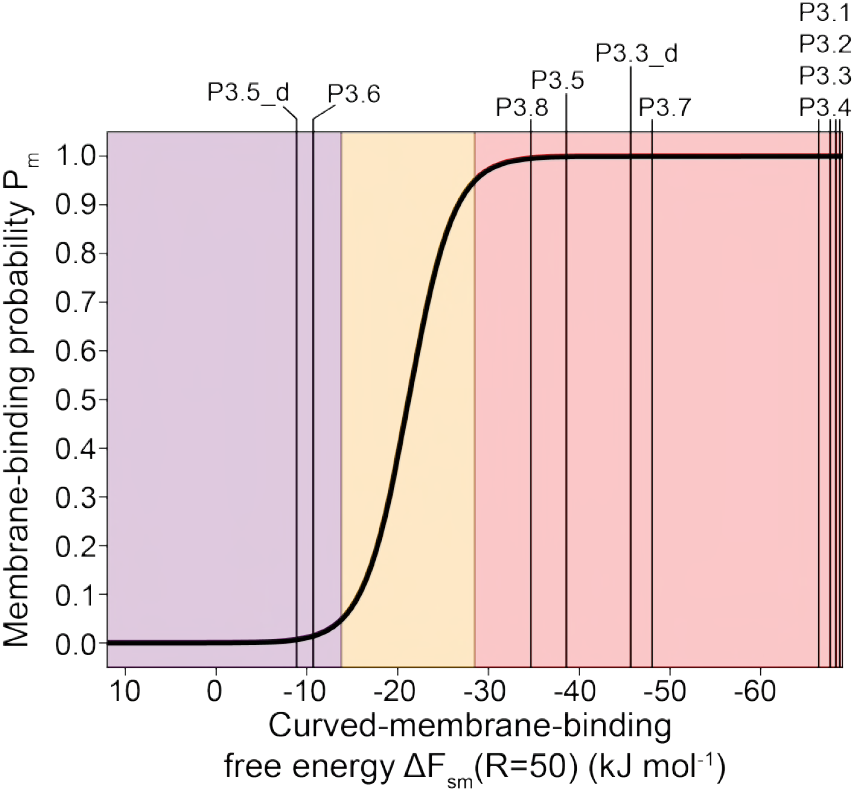
Predicted binding behavior of dual-objective optimized peptides. The membrane-binding probability *P*_m_ as a function of the curved-membrane-binding free energy, that is derived from the PMIpred-predicted ΔΔ*F*, as described in Ref. [25]. The dual-objective optimized peptides from generation 3 span a wide range of free energies.

**Fig. SI5.**
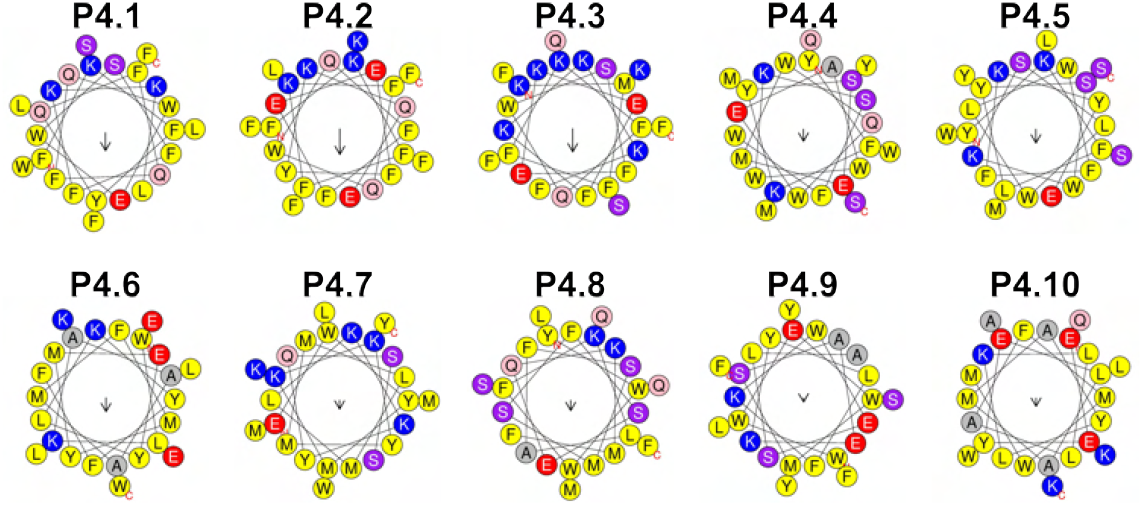
Helical wheels (generation 4). Helical wheel representations [37] for all P4.x peptides. Yellow: hydrophobic. Gray: small. Pink/purple: polar. Blue: positively charged. Red: negatively charged.

### 3 Experimental validation

#### 3.1 Generation P1.x and P2.x

##### 3.1.1 Antiviral activity and toxicity

To experimentally determine the antiviral activity of the Evo-MD-designed peptides, we incubated Zika virus (ZIKV) particles with increasing concentrations of peptide and then measured their infective potency on Vero E6 cells (see section SM3 for methodological details). ZIKV is a rather small virus (≈ 50 nm in diameter), and therefore it should be an ideal target for our peptides that are designed to bind to strongly curved membranes. Indeed, we found multiple peptides that potently inhibited ZIKV infection in our assay, with peptide P1.6 and P2.2.3 having the strongest effect (Fig. 2A). As a positive control, we measured HCV AH. As an example of an inactive Evo-MD-designed peptide, we also included P2.2. To determine toxicity, we incubated Vero E6 cells under the same condition but in the absence of virus. This showed that of the four highlighted peptides, only P2.2.3 was toxic at the used concentrations (Fig. e2H). Notably, P1.6 did not show any toxicity in this assay, being more than 71.4 times as selective in inhibiting ZIKV than disrupting Vero E6 cells (selectivity index SI = CC_50_/IC_50_ >71.4, see Table 1) [23]. For HCV AH, we determined the SI to be >16.7 for ZIKV/Vero E6 (Table 1). Overall, we found the cell viability data to be consistent with results from our additional hemolysis assay on erythrocytes (Fig. SI11, method is described in section SM3), again confirming no toxicity for P1.6 up to a 200 *µ*M concentration.

To probe the size-specificity of our peptides, we then performed a similar antiviral assay using a larger virus, HIV-1 (100-150 nm diameter, see section SM3 for methodological details). Here, we found that peptides that inhibited ZIKV typically also inhibited HIV-1 (Fig. SI6 and Fig. SI9), indicating that the mechanism of action is indeed membrane- and not protein-mediated.

**Fig. SI6.**
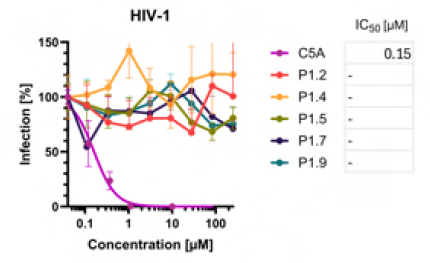
HIV-1 infectivity. Results of the generation P1.x against HIV-1 NL4-3 viruses that were not included in Ref. [23]

**Fig. SI7.**
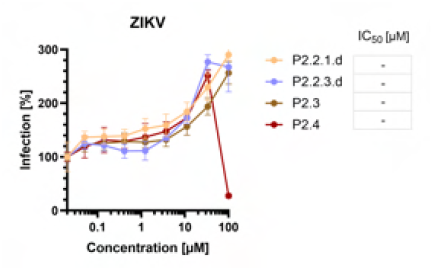
ZIKV infectivity. Results of the generation P2.x against ZIKV virus that were not included in Fig. 2A (N=1 in triplicates).

**Fig. SI8.**
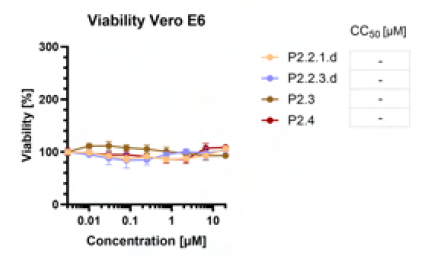
Vero E6 viability. Viability results for the generation P2.x against Vero E6 cells for the peptides that were not included in Fig. 2H (N=1 in triplicates).

**Fig. SI9.**
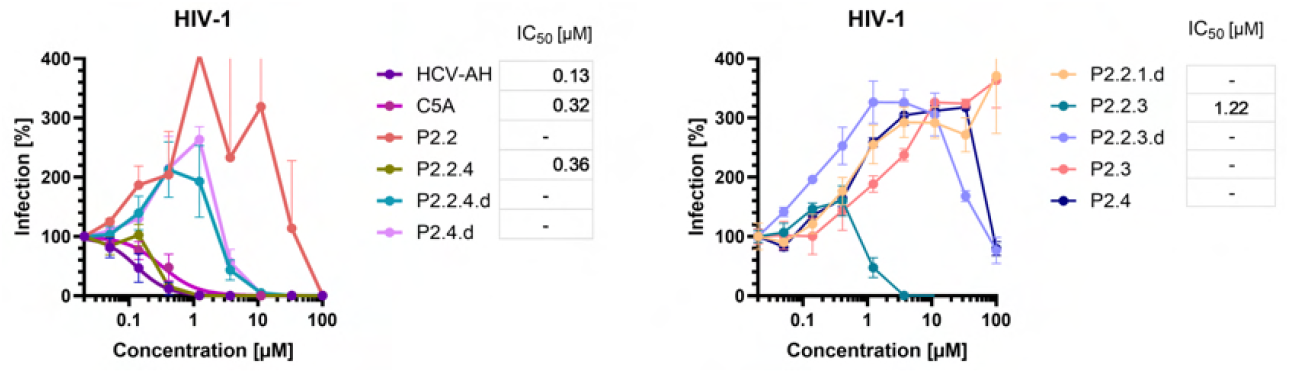
HIV-1 infectivity. Results for the generation P2.x against HIV-1 NL4-3 viruses (N=3 and N=1 in triplicates, for the left and right panel, respectively).

**Fig. SI10.**
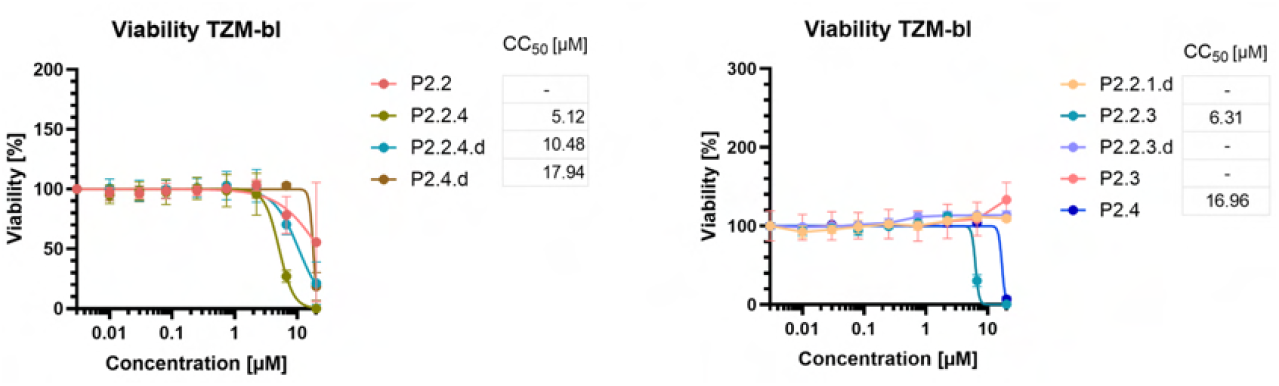
TZM-bl viability. Viability results for the generation P2.x against TZM-bl cells (N=2 and N=1 in triplicates, for the left and right panel, respectively).

**Fig. SI11.**
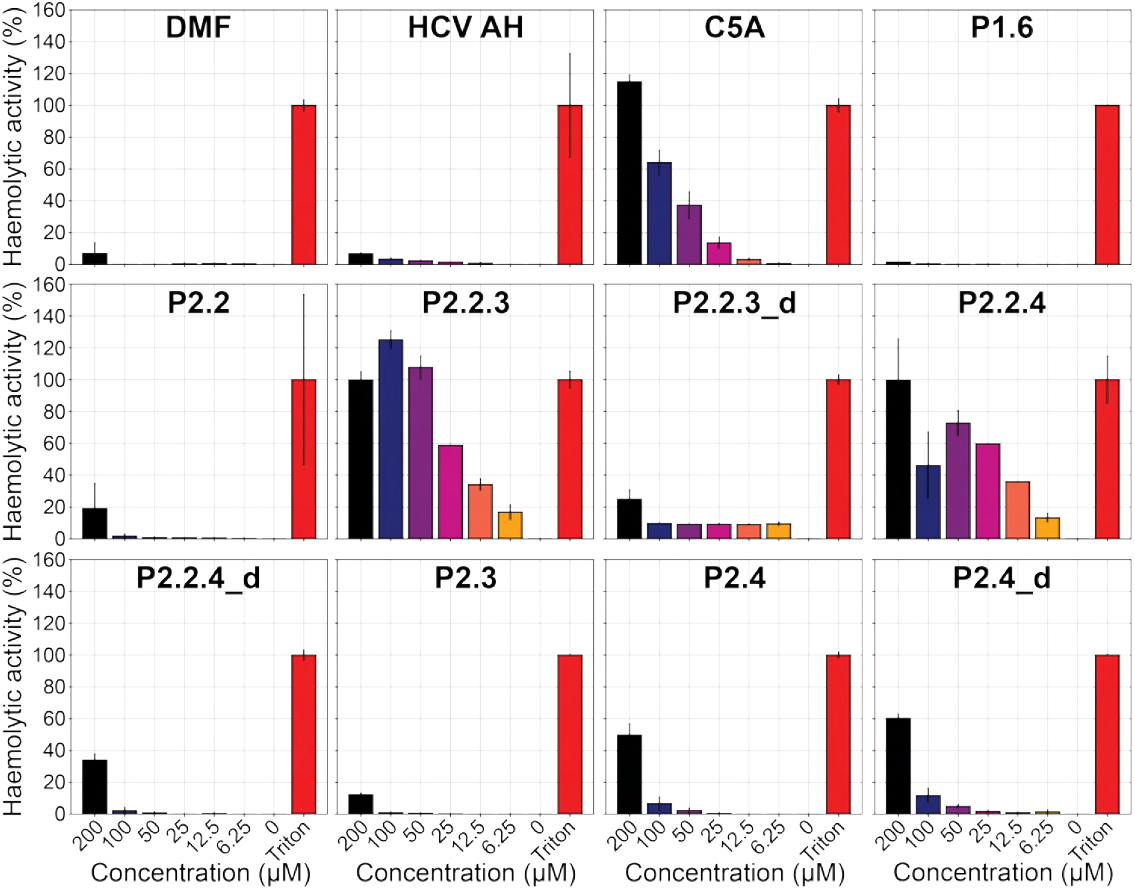
Hemolysis activity. Different concentrations of peptide (or DMF solvent) were added to erythrocytes that were isolated from fresh blood samples. Hemolytic activity was measured as described in section SM 3. Triton X-100 was added to determine the maximal activity (set to 100%).

Next, to test size-dependent activity in a more controlled biophysical setting, we employed a dye leakage assay [32]. In this experiment, liposomes of different sizes (50, 100, and 200 nm diameter) and a virus-like lipid composition (45/25/30 mol% DOPC/SM/cholesterol) with encapsulated fluorescein dye were prepared and then, at *t* = 0 min, incubated with increasing concentrations of peptide after which the dye leakage was measured. At *t* = 30 min, Triton X-100 was added to fully lyse the vesicles and establish maximal leakage (see section SM3 for methodological details). In line with the general trend in Fig. 2A, P1.6 most potently permeabilized the liposomes (EC_50_ = 0.05 *µ*M for 50 nm particles) [23], followed by P2.2.3 (a sequence derived from P2.2) and the positive control HCV AH (Fig. SI12). HCV AH was the only peptide for which we observed a slight size-dependent permeabilizing effect [23], as described previously [16, 17], although this was not reflected in the IC_50_ values on ZIKV (1.2 *µ*M, at a 50 nm diameter) and HIV-1 (0.1 *µ*M, at a 150 nm diameter) [23]. For the original peptide sequence P2.2, we did not observe any liposome leakage below 3 *µ*M concentration SI12, in agreement with its inability to inhibit ZIKV and HIV-1 infection. Additional adsorption experiments with DMPC monolayers showed that P2.2, in contrast to P1.6 and P2.2.3, does not integrate into the lipid environment, likely due to self-aggregation (Fig. SI13. Liposome leakage data for all remaining peptides are shown in Fig. SI12 and summarized in Table SI3.

**Fig. SI12.**
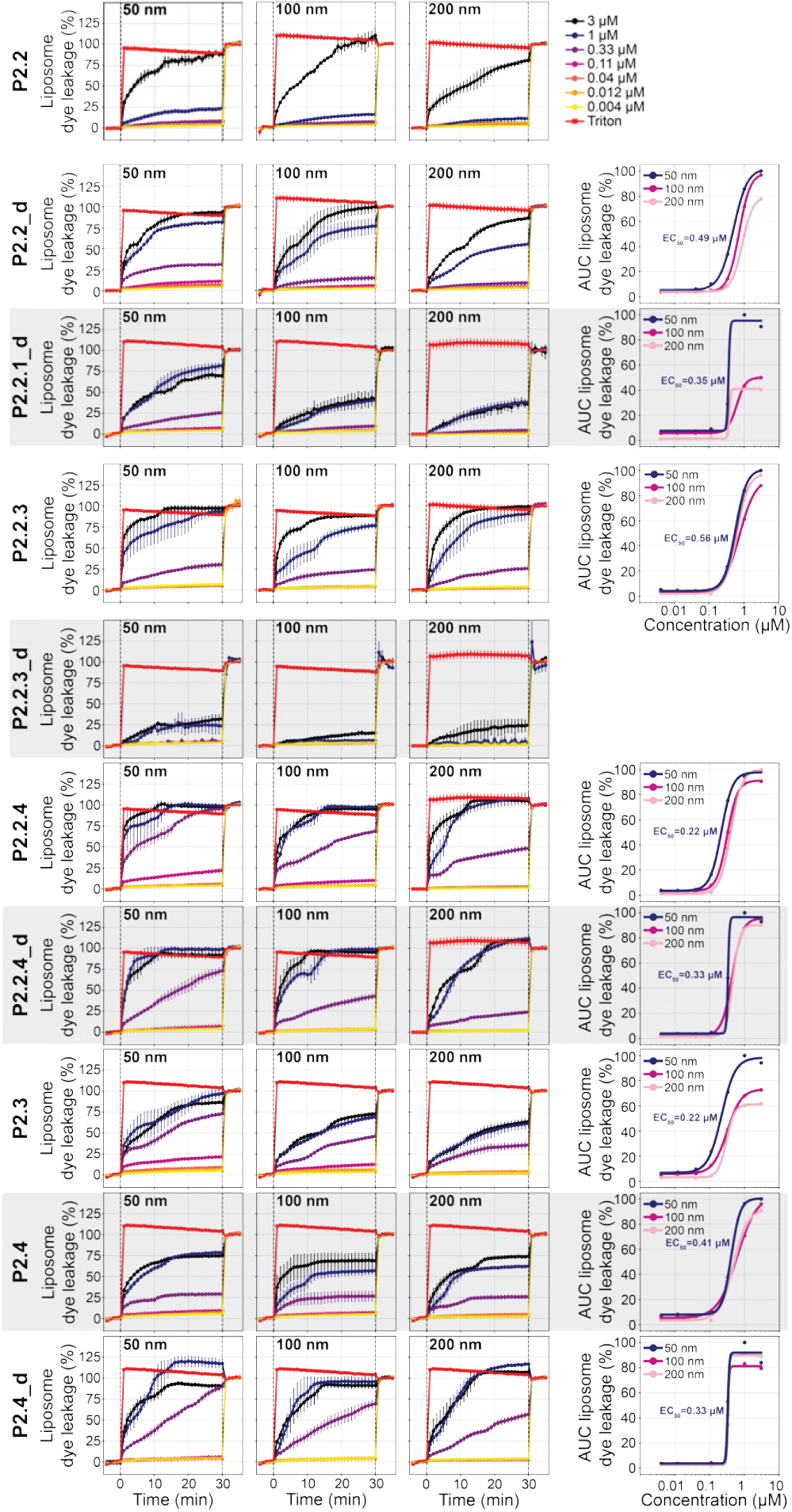
Size-dependent liposome leakage assay. Liposome dye leakage from 50, 100, and 200 nm model liposomes (45/25/30 mol% DOPC/SM/cholesterol) [32] upon addition (at *t* = 0 min) of different concentrations of peptide, or Triton X-100. At *t* = 30 min, Triton X-100 was added to determine the maximal leakage (set to 100%). In the final column, we plotted the area-under-the-curve (AUC) at different liposomes sizes; only shown if leakage exceeded 50% for at least 2 peptide concentrations for any liposome size. EC_50_ values were derived from sigmoid fits. The inset EC_50_ values are calculated for the 50 nm liposome data.

**Fig. SI13.**
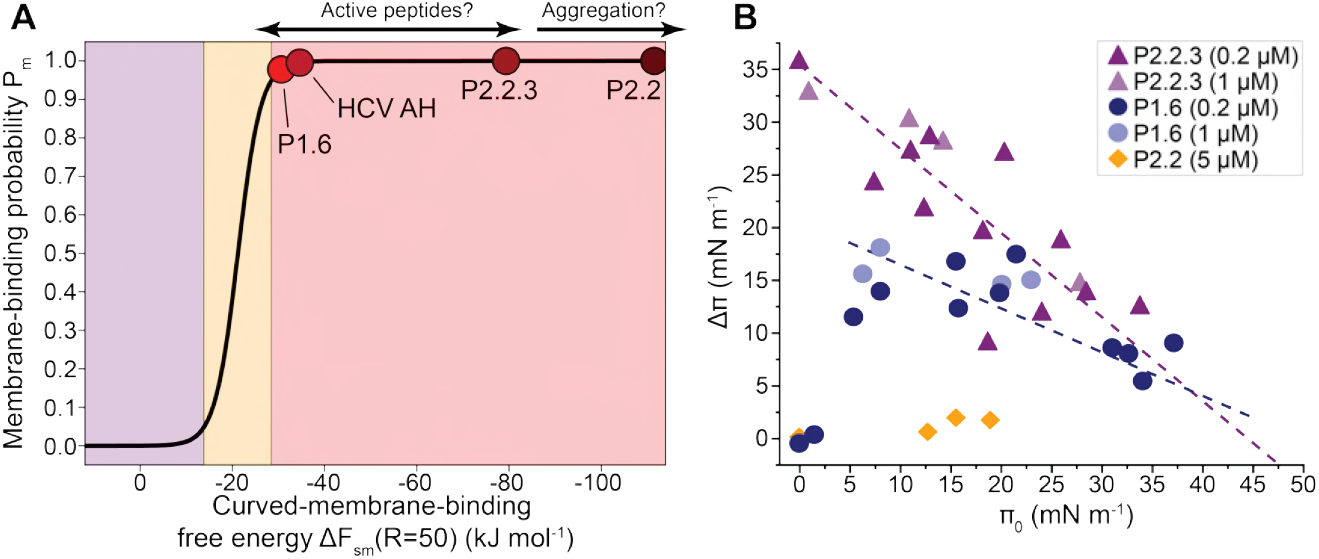
Membrane-binding ability of active peptides. **A)** Using the thermodynamic model from Ref. [9], we find that P1.6, HCV AH, and P2.2.3 define a lower binding regime where active peptides are located. P2.2 is so “binding-prone” that it may self-aggregate and not bind to the membrane at all. **B)** Interaction of peptides with DMPC monolayers (25°C; 150 mM NaCl; pH 7.4). The change in surface pressure Δ*π* after injection of the peptide into the subphase of preformed DMPC monolayers is depicted as a function of initial surface pressure *π*_0_. The intercept Δ*π* = 0 yields the maximal insertion pressure (*π*_mip_) which can be related to the ability of the peptides to insert into bilayers. Data points with *π*_0_ below 2.5 mN m^−1^ were obtained without lipid monolayer, i.e. they represent the interaction of the peptide with the air-buffer interface.

##### 3.1.2 Pore stabilization in MD simulations

To provide some molecular insight into how our Evo-MD-designed AVPs may disrupt membranes, we performed coarse-grained MD simulations of peptide-assemblies in a perforated membrane. The starting configuration of these simulations was a 6-mer bundle of peptides assembled inside a membrane pore that was externally enforced by a cylindrical flat-bottom (FB) potential acting on the lipid tails (Fig. SI14A). We observed that the presence of all three peptides P1.6, P2.2.3, and HCV AH reduced the FB potential energy required to keep the pore open (although not statistically significant for HCV AH, see Fig. SI14B). Next, we asked what would happen once we switched off the FB potential (Fig. SI14A): would the pores be stable? Or would they disintegrate or collapse? Interestingly, we observed that – for both our Evo-MD-designed AVPs P1.6 and P2.2.3, and the literature standard HCV AH – the pores shrunk but retained a stable water channel during our 2 *µ*s simulations, as evident from the non-zero solvent densities in Fig. SI14C. We also observed that in all cases, some peptides were expelled from the initial 6-mer bundle after the FB potential was released, although the stoichiometry of the final pore structures varied between the three different peptides (Fig. SI14D). We found that P1.6 formed 4/5-mer pores, with three peptides being fully transmembrane and two only contributing transiently. HCV AH formed a more classical 4-mer toroidal pore structure with all four peptides adopting a tilted conformation. Finally, P2.2.3 formed an asymmetric 2-mer pore, possibly indicating a different disruptive mechanism from the other two peptides tested.

**Fig. SI14.**
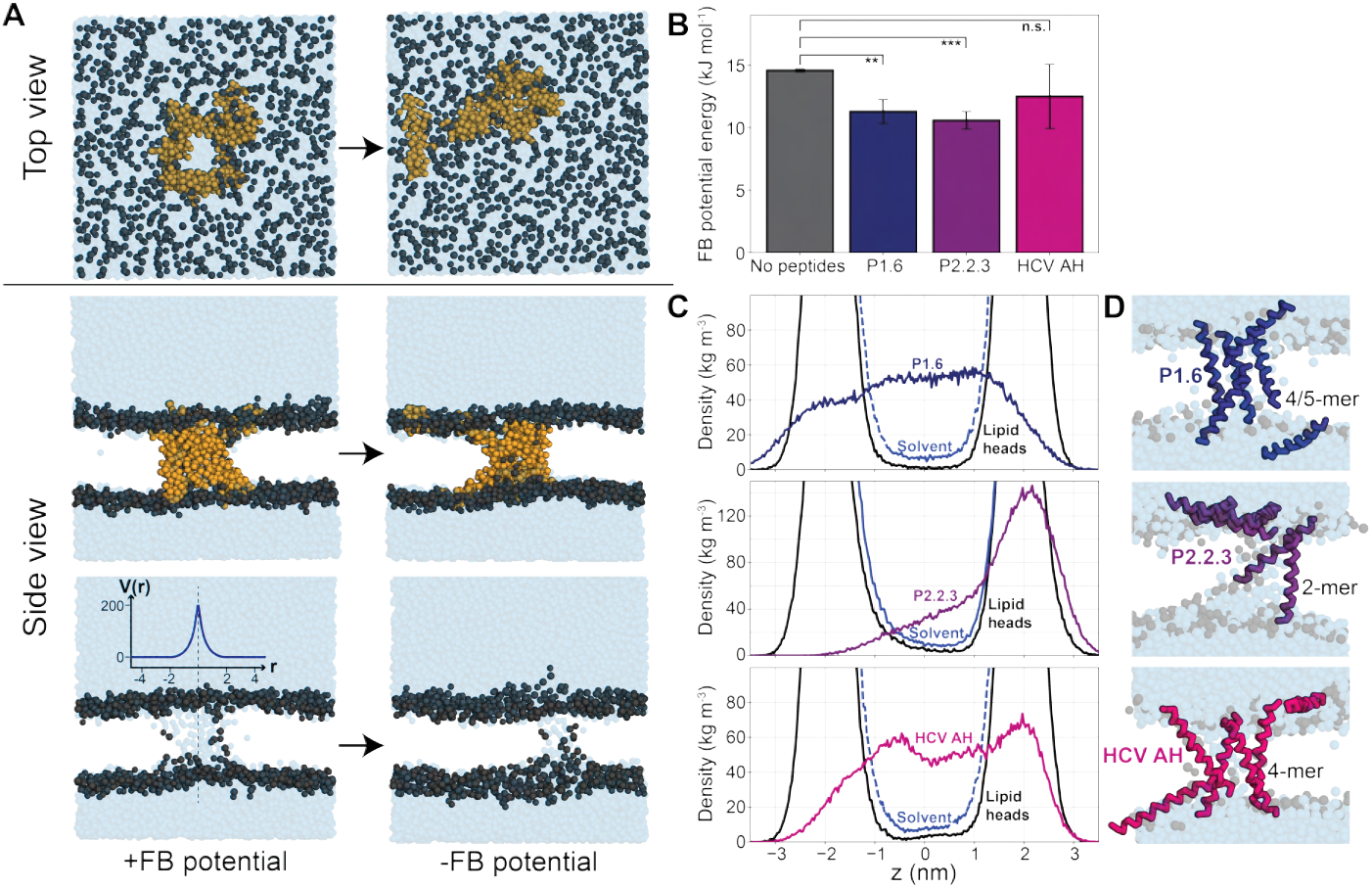
Antiviral peptides stabilize pores in MD simulations. **A)** Top view and side view of the initial 6-mer bundle setup while applying the pore-forming FB potential (left) and the final configuration, without the FB potential (right). Solvent is shown in blue. POPC head groups are shown in gray. POPC tails are hidden for clarity. The example peptide (in orange) is HCV AH. **B)** Peptide bundles reduce the FB potential energy required to keep the pores open. Means and standard deviations from ensemble averaged values of the final 500 ns from 5 independent rerun trajectories are shown. P-values were calculated with the two-tailed Welch’s t-test (^*^*p* < 0.05, ^**^*p* < 0.005, ^***^*p* < 0.0005). **C)** Trajectory-averaged density plots along the *z*-axis of the three different peptides, solvent (water and ions) and POPC head group beads (NC_3_ and PO_4_), centered on the membrane *xy*-midplane. The first 500 ns were discarded for equilibration. **D)** Representative MD snapshots of the final peptide assemblies that stabilize the membrane pore. Solvent is shown in blue. POPC head groups are shown in gray. POPC tails are hidden for clarity. Panel A and D were adapted from Ref. [23].

##### 3.1.3 Assembly and methodological details of pore stabilization simulations

We prepared and equilibrated a Martini 3 POPC membrane with 225 lipid molecules in each leaflet and with 0.15 M of Na^+^ and Cl^−^ ions added to the solution (settings conform with Ref. [24]). Then, we applied an inverted cylindrical flat bottom (FB) potential 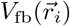 that only acts on the POPC tail beads (*i*) and uses the *z*-axis through the center of the membrane as the reference coordinate 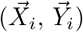:

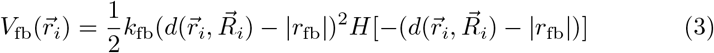

in which 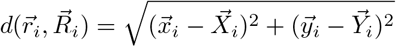 and *H* is the Heaviside step function (see the GROMACS user manual for details [60]). For our simulations, we chose *k*_fb_ = 100 kJ mol^−1^ and *r*_fb_ = −2 nm to get the potential shape plotted in the insets in Fig. SI14A and Fig. SI15A. Due to instabilities caused by the external FB potential, we reduced the step size to 10 fs in all corresponding simulations. 6-mer peptide bundles in the pore were obtained through self-assembly from an initial setup with 6 peptides in a 3 × 2 grid in an *xy*-plane, 5 nm from the membrane surface (Fig. SI15A). Within 2 *µ*s of simulation, P1.6 and HCV AH readily formed a 6-mer toroidal pore structure by adhering to the strongly curved edges of the pore rim, as evident from the *z*-distance trajectories (five independent reruns, see Fig. SI15B). Conversely, the six P2.2.3 peptides aggregated in the water phase and did not assemble in the pore (middle panel in Fig. SI15B). To probe if P2.2.3 could assemble into a toroidal pore structure at lower concentrations, we added the peptides one by one at 2 *µ*s intervals and observed that the individual peptides indeed readily translocate to the pore rim (Fig. SI15C), although only four peptide molecules were positioned inside the pore in the final configuration (at 12 *µ*s in Fig. SI15C).

The final self-assembled structures (at 2 *µ*s in Fig. SI15B for P1.6 and HCV AH and at 12 *µ*s in Fig. SI15C for P2.2.3) were used as the starting topologies for the simulations in Fig. SI14.

**Fig. SI15.**
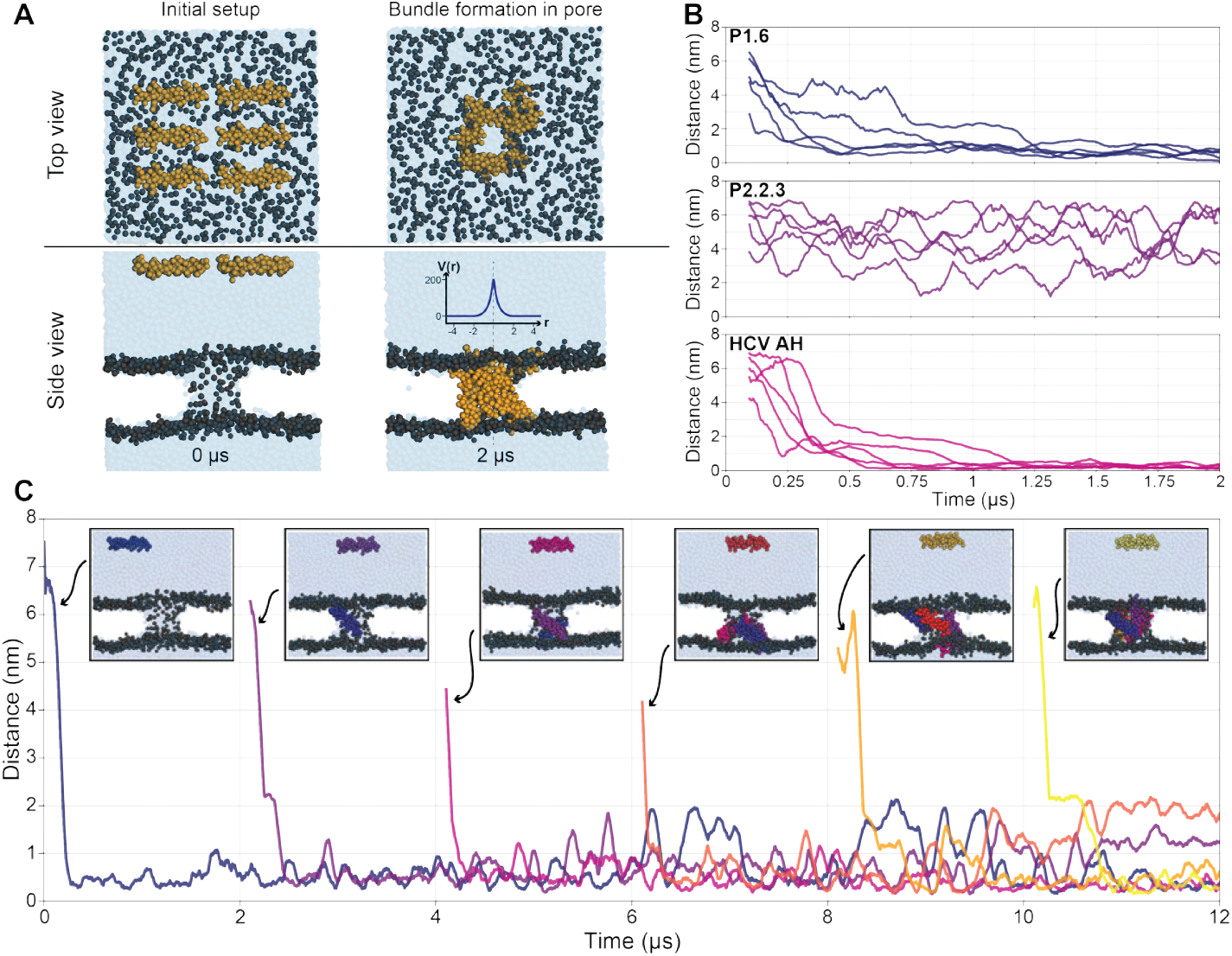
Self-assembly of peptides in a membrane pore. **A)** Top view and side view of the initial simulation setup (left) and the self-assembled 6-mer bundle in the membrane pore after 2 *µ*s (right). The inset in the bottom right panel shows the shape of the inverted FB potential, eq. 3. **B)** *z*-component of the distance between the center-of-mass of the six peptides and the center plane of the membrane over five independent reruns for peptides P1.6, P2.2.3, and HCV AH (100 ns running averages). **C)** *z*-distance between the center-of-mass of the individual P2.2.3 peptides and the membrane center plane (100 ns running averages). Insets show starting configurations upon addition of the next peptide (every 2 *µ*s).

#### 3.2 Generation P3.x: ZIKV and HIV-1 inhibition data

**Fig. SI16.**
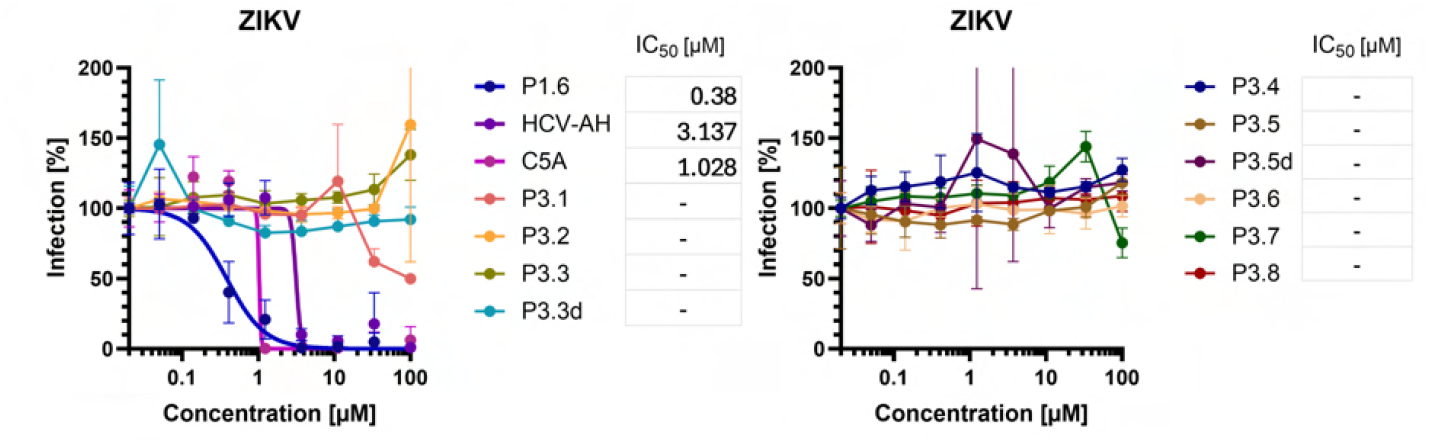
ZIKV infectivity. Results of the generation P3.x against ZIKV viruses (N=1 in triplicates). No significant antiviral activity was observed

**Fig. SI17.**
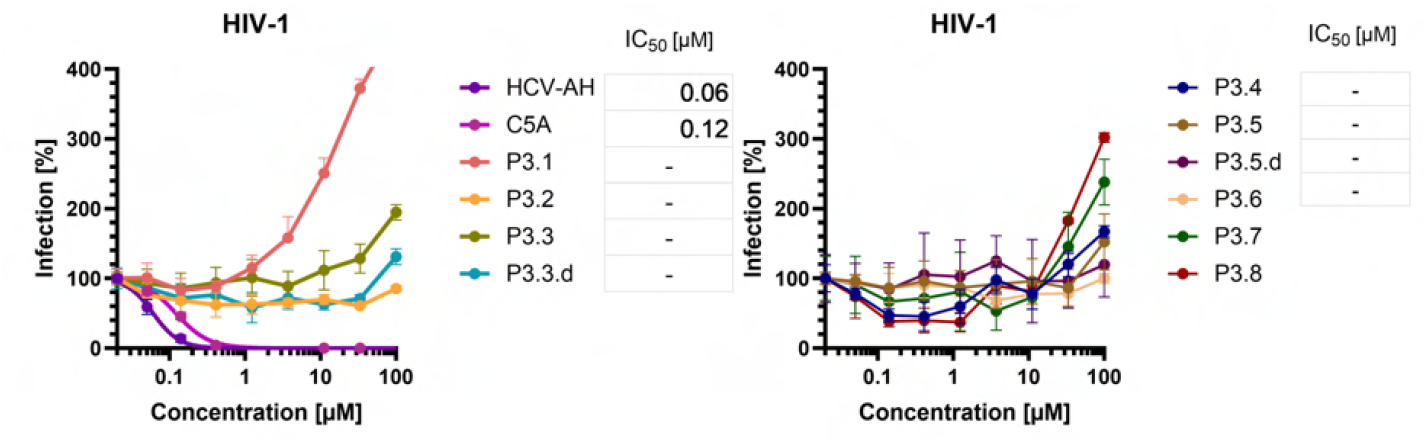
HIV-1 infectivity. Results of the generation P3.x against HIV-1 NL4-3 viruses (N=1 in triplicates). No significant antiviral activity was observed.

#### 3.3 Generation P4.x: ZIKV and HIV-1 inhibition data

**Fig. SI18.**
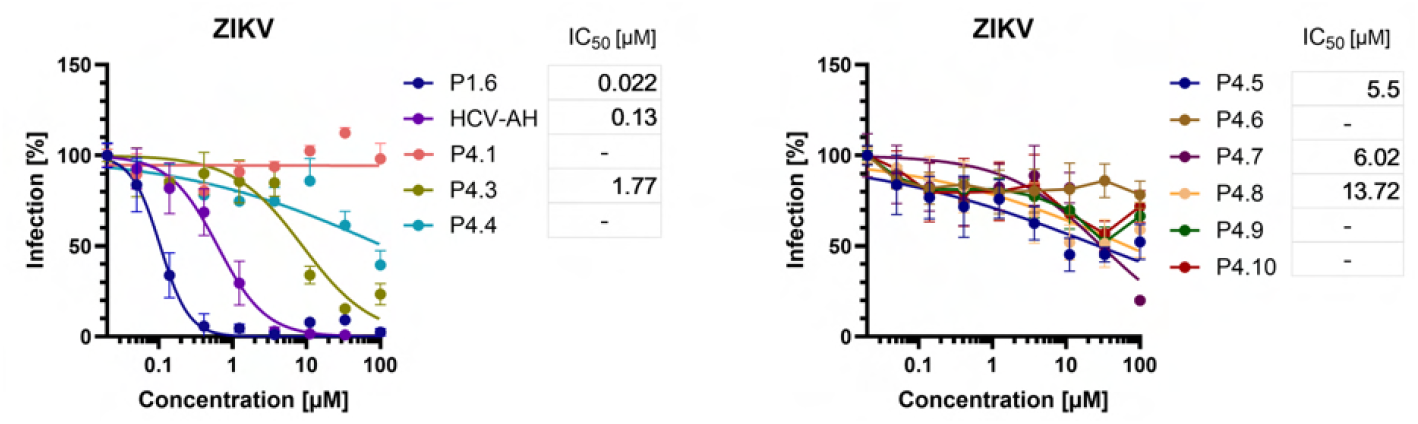
ZIKV infectivity. Results of the generation P4.x against ZIKV viruses (N=1 in triplicates).

**Fig. SI19.**
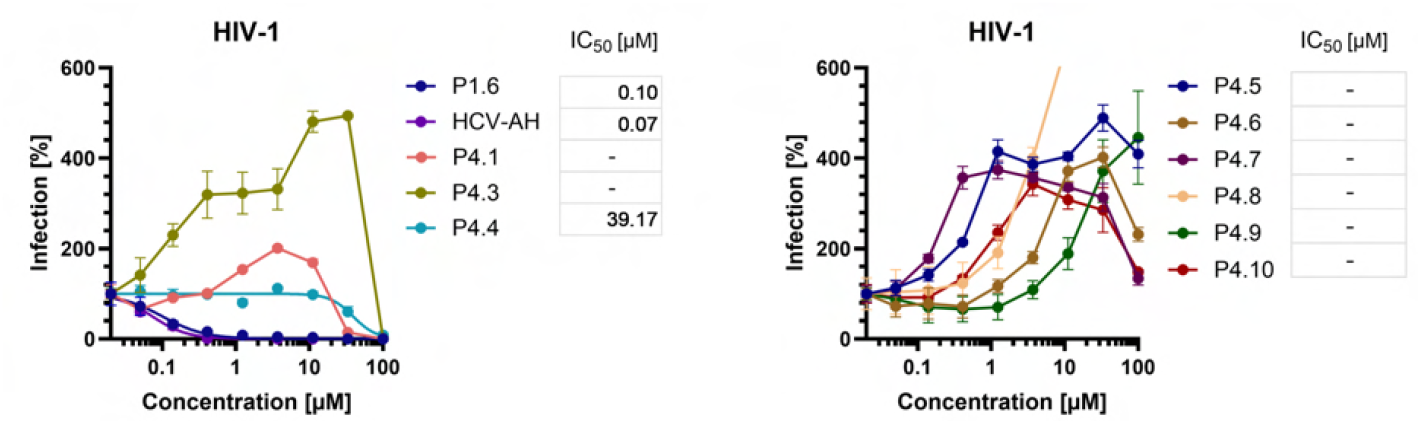
HIV1 infectivity. Results of the generation P4.x against HIV1 viruses (N=1 in triplicates).

**Fig. SI20.**
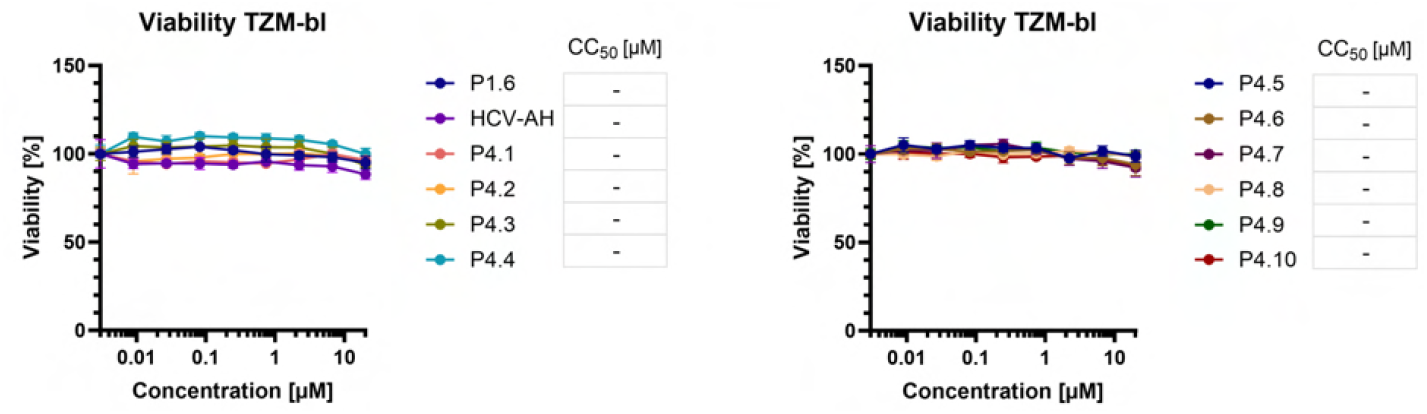
TZM-bl viability. Viability results for the generation P4.x against TZM-bl cells (N=1 in triplicates).

